# Ageing via sexual perception is a by-product of male adaptive plasticity in *Drosophila melanogaster*

**DOI:** 10.1101/2021.04.19.440494

**Authors:** Quentin Corbel, Manuel Serra, Roberto García-Roa, Pau Carazo

## Abstract

Sensory perception of environmental cues can dramatically modulate ageing across distant taxa. For example, male *Drosophila melanogaster* age faster if they perceive female cues but fail to mate (ageing via sexual perception). This finding has been a breakthrough for our understanding of the mechanisms of ageing, yet we ignore how and why such responses have evolved. Here, we used *D. melanogaster* to ask whether ageing via sexual perception may be a by-product of plastic adaptive responses to female cues, and found that while long-term sexual perception leads to reproductive costs, short-term perception increases male lifetime reproductive success in a competitive environment. Simulations under a wide range of socio-sexual and demographic scenario suggest that such plasticity as a response to sexual perception might be a widespread strategy in nature. Finally, we show that sexual perception can significantly magnify sexual selection (15-27% average increase in the opportunity for selection).

## INTRODUCTION

Over the last decades, we have realised the potential for sensory perception to act as a modulator of ageing across invertebrate and vertebrate taxa (1–4). A series of ground-breaking studies have shown that sensory perception can trigger physiological changes at multiple levels, from homeostasis to tissue physiology, that significantly accelerate ageing (3, 5–8). For instance, chemosensory perception of food impairs the extended lifespan conferred by dietary restriction in both the vinegar fly *Drosophila melanogaster* (3) and in the nematode *Caenorhabditis elegans* (1, 9). Similarly, in these two model species, exposure to different types of conspecific cues decreases lifespan and alters critical physiological traits (5,7,8,10), suggesting that ageing via sensory perception operates across distant taxa and functional contexts.

Organisms respond to environmental cues through a host of plastic physiological, anatomical or morphological changes that allow them to adjust their behaviour and life-history strategy so as to improve their fitness (11). For example, in some insect species, males respond to socio-sexual cues (i.e. density and diversity of rival male odours indicative of varying levels of sperm competition) by strategically adjusting their investment in sperm numbers, ejaculate components and/or post-copulatory mate guarding behaviour (e.g. 12-16). Plastic responses are also sometimes maladaptive and lead to a decrease in individual fitness. For example, stressful conditions or unreliable environmental cues can sometimes engender maladaptive plasticity (17, 18) and can possibly hinder the evolution of adaptive genotypes (19–20). Thus, in principle, ageing via sensory perception may result from either adaptive or maladaptive responses to environmental cues. Because studies have so far primarily focused on identifying the mechanisms linking sensory perception to ageing, we simply ignore what selective pressures (if any) underlie this phenomenon. It has been suggested that long-term costs associated with sensory perception could be a by-product of adaptive responses in the short-term (21), but the question of whether ageing via sensory perception bears an adaptive value remains unanswered.

Sexual perception (i.e. perception of reproductive opportunities) is a particularly interesting driver of ageing. *Drosophila melanogaster* males that do not regularly mate suffer significant physiological costs and a drastic reduction in survival as a sole consequence of perceiving female chemical cues (5, 7). In addition, long-term exposure to female chemical cues reduces male lifetime fitness in a competitive environment by approximately 25%, mostly due to accelerated reproductive ageing (22). Given that nature showcases a wide range of sophisticated adaptive plastic responses to diverse and complex types of reproductive cues, understanding why sexual perception accelerates ageing and whether it bears an adaptive value may be of particular evolutionary interest. In addition, it has been suggested that ageing via sexual perception can magnify sexual selection (22, 23). Because ageing via sexual perception is contingent on a failure to mate, it will act to further decrease the fitness of males with inherently low mating success, thus potentially increasing the overall variance in male reproductive success (i.e. the opportunity for selection) and consequently the potential for sexual selection. Furthermore, if costs associated with long-term sexual perception are a by-product of adaptive short-term responses, sexual perception may similarly intensify sexual selection by increasing the inherent fitness advantage of high-quality (i.e. high probability of mating soon and acquiring sensory-mediated benefits) vs. low-quality (i.e. low probability of mating soon) males. Sexual selection is pivotal in driving male and female fitness, population viability, and evolvability (24–31), hence the need to address the potential role of ageing via sexual perception as a magnifier of sexual selection (23).

In this study, we first address the question of whether ageing via sensory perception can be adaptive by disentangling the fitness consequences of short vs. long-term sexual perception in *D. melanogaster*. We individually exposed males to females (treated males) while preventing access to females (experimentally simulating repeated failure to mate) for a period of 1, 3, 7 or 15 days, thus artificially manipulating the onset of mating. We then compared the lifetime reproductive success of treated vs. controls when competing for a female against two rival males and found that male lifetime fitness increased after a short-term exposure (1 day) to female cues, was neutral following an intermediate exposure time (3 and 7 days), and only resulted in lower lifetime fitness following long-term exposure (15 days) to female cues. Rate-sensitive fitness analyses highlighted the importance of late-life reproduction (see 32,33) at modulating these effects, implying that both beneficial and costly plastic responses acquired through sensory perception are long-lasting. Next, we asked under what mating systems and demographic scenarios we may expect such male plasticity to evolve. We used the empirical results obtained to parameterize mathematical models and ran simulations under a wide range of socio-sexual contexts varying in mating rate (promiscuity), variability in male mating success (i.e. opportunity for sexual selection) and background population growth (i.e. demography). We found average mating rate to be the critical component for male plasticity to evolve, such that male plastic responses to female cues are favoured in promiscuous populations/species. Finally, we used these same simulations to estimate the degree to which male plastic responses in response to sexual perception may magnify sexual selection, and found that the opportunity for selection can be increased significantly (i.e. 15 to 27% average increase).

## RESULTS

### Reproductive success

While short-term sexual perception improved reproductive performances of males (vs. controls), longer exposures lead to impaired reproductive performances, relative to control flies (i.e. significant exposure x treatment interaction in a LM: *F_1,405_=* 4.81, *P*= 0.029; Figure 1). A population growth rate-sensitive analysis of fitness yielded qualitatively similar results, with a clear fitness advantage in the treatment group after 1 day of sexual perception across all population growth rates, but particularly so in a declining population (Figure 2a). This advantage mostly disappeared following 3 days and 7 days of sexual perception, except under markedly negative population growth rate (Figure 2bc). Finally, longer exposures of 15 days and 21 days (21 days data taken from 22) showed clear fitness costs across all population growth rates, even more so in a declining population context (Figure 2de). Generally, the magnitude of the interaction between exposure time and treatment was strongest in a decreasing population demographic scenario (relative to stable and growing population) and weakened with increasing growth rates (Figure 2; see also SM).

**Figure 1:**
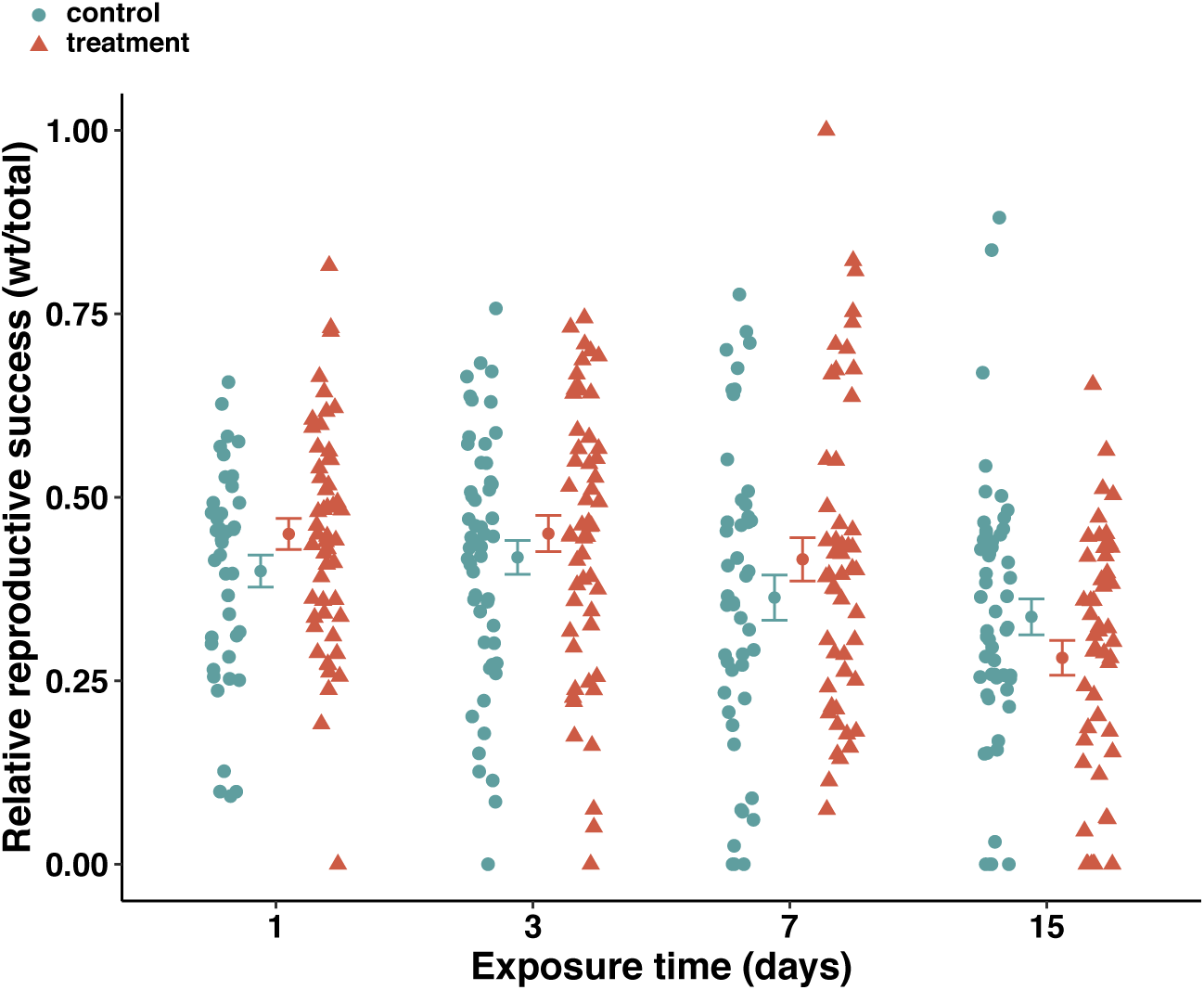
Relative lifetime reproductive success (percentage of offspring fathered by the focal male) after 1 day, 3 days, 7 days and 15 days exposure to the sensory treatment or control. Orange triangles represent treated males and green circles represent control males. Inwards these single observations are displayed means ± SEM. To avoid superposition of the datapoint belonging to different experimental conditions, control and treatment datapoints of the same exposure were presented separated from each other on continuous x axis.

**Figure 2:**
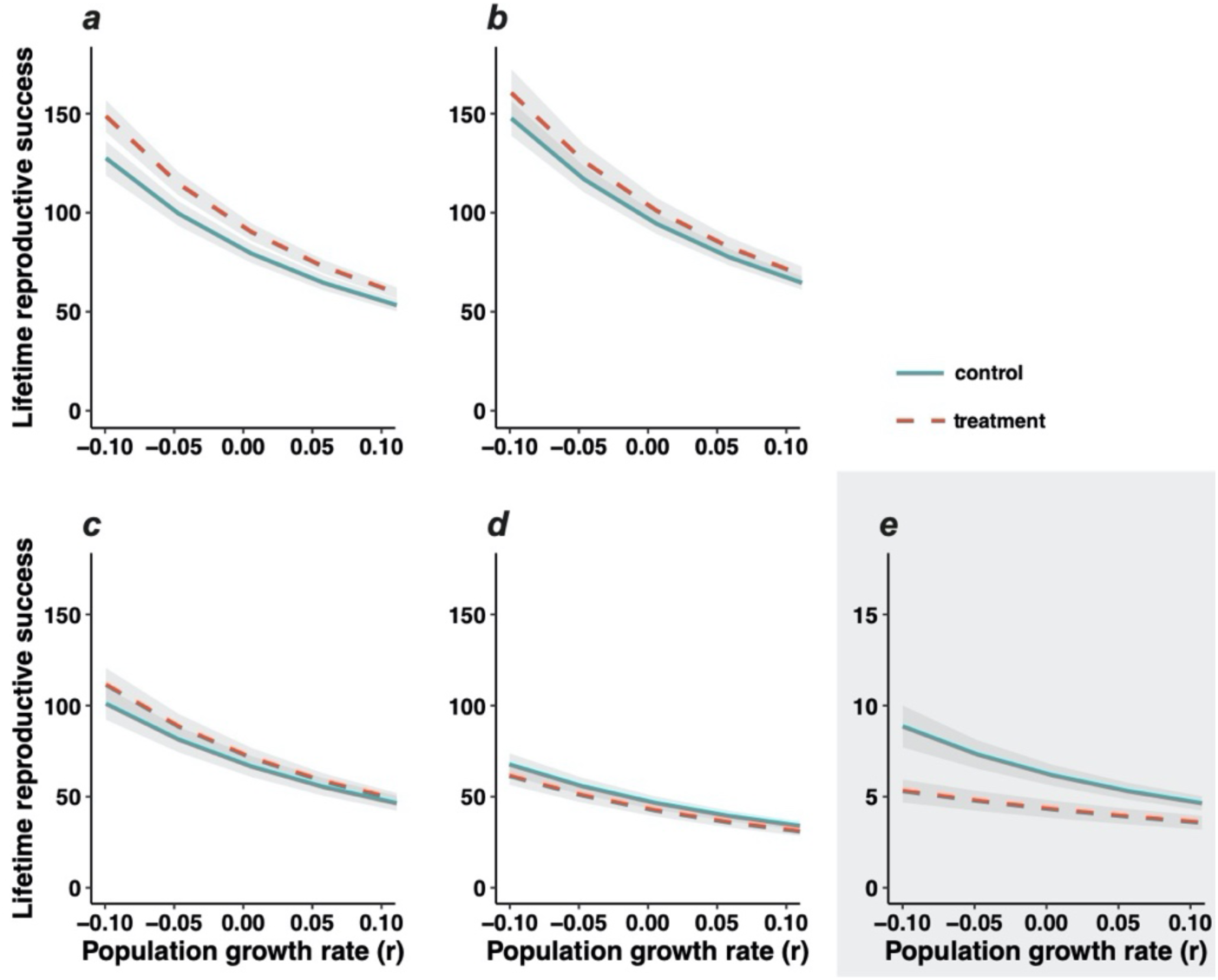
Average lifetime reproductive success of treated (red dashed line) versus control (green continuous line) males as a function of population growth rate, after 1 day, 3 days, 7 days, 15 days and 21 days of exposure to female conspecifics (***a****, **b**, **c**, **d**, **e**,* respectively; ***e*** after García-Roa *et al.* (22)). Lines are linear interpolations between means computed for each population growth rate. The greyed area represents the standard error around the mean. ***e*** is included to provide context. Because flies were older when provided access to mating, average fitness is considerably lower, regardless of the treatment. Therefore, the response variable (lifetime reproductive success) is downscaled by 10.

### Survival

A Cox proportional hazards survival model revealed a significant exposure x treatment interaction (LR*χ^2^_1_* = 5.14, *P*= 0.023).

### Reproductive lifespan and reproductive ageing

We did not find significant effects on either reproductive lifespan (i.e. treatment x exposure effect, LR *χ^2^_1_* = 0.09, *P*= 0.767; sensory treatment effect, LR *χ^2^_1_* = 0.35, *P*= 0.555; exposure time effect, LR *χ^2^ _1_*= 0.145, *P*= 0.704) or reproductive ageing (using net decrease in male relative reproductive success between the first and the second female they competed over –henceforth Δ♀_1_♀_2_–, *F_1,393_* = 0.22, *P*= 0.636; using net decrease in male relative reproductive success between the first and the third female they competed over – henceforth Δ♀_1_♀_3_–, *F_1,359_* = 1.38, *P*= 0.240). Sensory treatment had no effect on reproductive ageing (using Δ♀_1_♀_2_: *F_1,393_* =0.33, *P*=0.569; using Δ♀_1_♀_3_: *F_1,359_* = 2.00, *P*= 0.158), but exposure time significantly affected reproductive ageing (Δ♀_1_♀_2_: *F_1,393_* =7.56, *P*= 0.006; using Δ♀_1_♀_3_: *F_1,359_* =8.80, *P*= 0.003) as the decrease in relative reproductive success over the course of two (and three) females was of higher magnitude with increasing exposure (see SM for further details).

### Model and simulations

We built a computer model to estimate male fitness based on the assumptions that (1) the log-reproductive function (l(x) m(x); standard demographic notation) modelling realized reproductive lifespan (i.e. from the onset of mating (β) onwards) is a 90-degree rotated sigmoid curve (i.e. monotonously decreasing, levelling-off at intermediate ages) and that (2) the elevation (E) of that curve declines linearly with β. The features and additional details of our computer model were largely based on the fitting of ad hoc Generalized Additive Models to our empirical results. Another assumption of the computer model is that (3) the relationship between E vs. β differs according to two alternative strategies: a) a “spendthrift” strategy (henceforth *S* strategy), characterised by males that always respond as if perceiving female cues (i.e. always engage the physiological responses associated with perception of reproductive opportunities), and b) a “thrifty” strategy (henceforth *T* strategy), characterised males that never respond to female cues (i.e. never engage the physiological responses associated with perception of reproductive opportunities; see SM for further detail). Note that presence/absence of female cues is irrelevant to both these strategies, as they are unconditional fixed strategies. Our aim with these simulations was to explore what conditions favour the expression of the *S* fixed phenotype vs. the *T* fixed phenotypes vs. the evolution of a plastic phenotype that could display either of *S* or *T*. In our simulations, a randomly sampled β was assigned to each individual, and this allowed us to compute the effects of the *S* and *T* strategies on within-male population variation in fitness. We explored the *S* and *T* strategies in scenarios resulting from the combination of three factors: (1) daily population growth rate (r: −0.1, 0, 0.1), (2) mean value of β (from 1 to 15 d), and (3) standard deviation of β (σ_β_: 0.01, 0.1, 1d). We envision mean β as a direct consequence of female density, and we assume standard deviation of β to be correlated to the heritable variation in male mating performance. The effect of the mean value of β on simulated fitness followed a pattern tightly consistent with variation in E (Figure 3). This points out the prevalent role of E in mediating the effect of β on fitness. Our simulations showed that the *S* strategy was advantageous (compared to the *T* strategy) whenever mean β was lower than 7-8 days, and disadvantageous for higher mean β (Figure 3). More precisely, within a range of β between 1 to 7-8 days, the relative advantage of the *S* strategy increased with decreasing mean β. Contrastingly, within a range of β between 8 to 15 days, the disadvantage the *S* strategy increased with increasing mean β. Standard deviation in male fitness declined steeply with mean β with the *S* strategy, and was rather independent of mean β with the *T* strategy. This results in the *S* strategy leading to higher fitness variation than the *T* strategy for mean β lower than 7-8 days, but causing lower fitness variation for mean β higher than 7-8 days. For example, in a stationary population (r = 0), fitness was higher for the *T* strategy (38% in average) across average mean β equal or above 7 days. Contrastingly fitness was lower for the *T* strategy (35% in average) across average mean β below 7 days. With respect to the increase in male fitness variation (i.e. opportunity for selection), we found that responding to female cues (i.e. plastic male strategy switching between the *S* and the *T* strategies accordingly to what strategy is advantageous) resulted in an overall average increase in standard deviation of 15 to 27%. Within-population standard variation in onset of mating (male mating rate; i.e. a proxy for variation in male quality and hence starting opportunity for selection) had little effect on mean fitness and standard deviation of fitness, so estimates above are given for the median across the whole range explored (see SM). These patterns are qualitatively the same for the population growth rates and the standard deviation of β explored in all simulations (see SM for further details).

**Figure 3.**
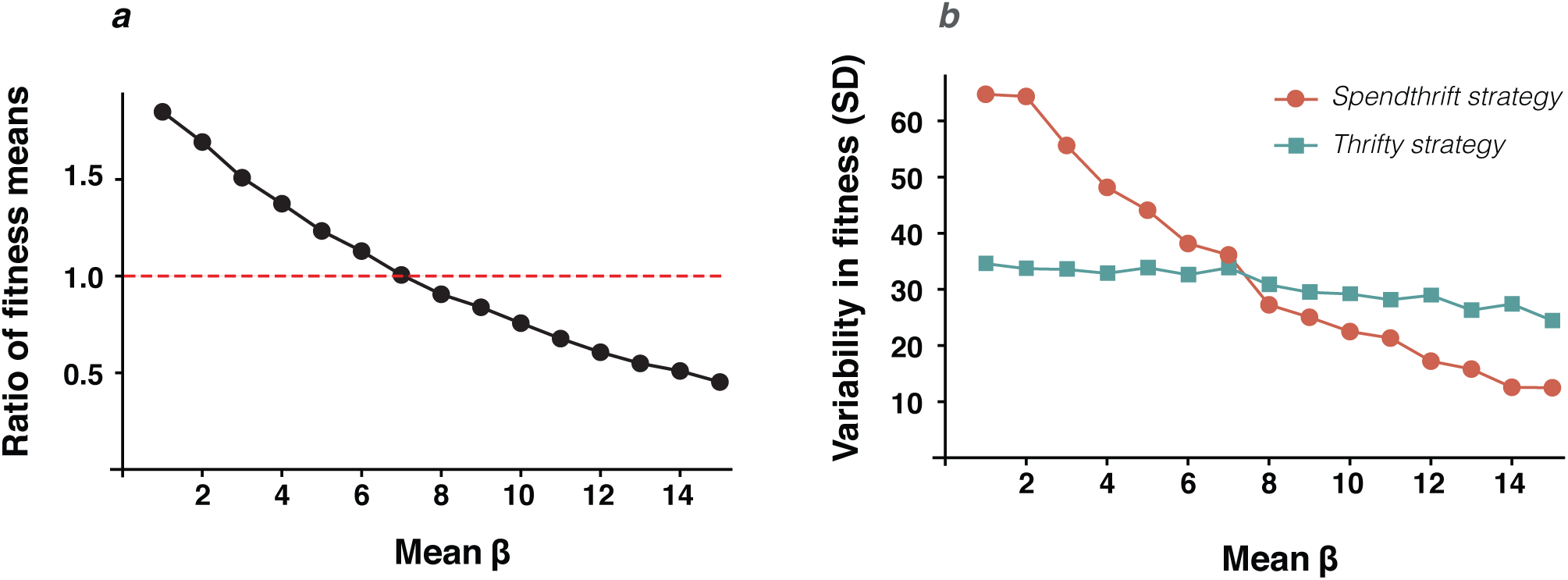
Ratio between male fitness means (**a**) and male fitness standard deviations (**b**) for simulated fly males across values of mean β (age of mating onset). Here the simulation portrays a growing population (r = 0.1) with standard deviation of β = 1 and sample size n= 10000. In (**a**), y values above one represents an advantage of the spendthrift strategy over the thrifty strategy, whereas values below one represent a disadvantage of the spendthrift strategy over the thrifty strategy.

## DISCUSSION

In this study, we found evidence strongly suggesting that the so far reported ageing effects induced by male perception of female cues in fact derive from adaptive male plastic responses. Sensing female cues for a short period of time (1 day) prior to reproduction triggered plastic responses in males that made them better competitors, and ultimately increased their lifetime reproductive success. Perception of female cues was neutral following intermediate exposures (between 3 and 7 days) and only resulted in net fitness costs if males perceived females for at least 15 days of simulated reproductive failure. Simulations showed that such plastic response can be advantageous under a wide range of socio-sexual and demographic contexts and significantly increase the opportunity for selection, and thus the intensity of sexual selection.

We found that, in *Drosophila melanogaster*, short-term (1 day) sensory exposure to female cues (simulating a short lag between perception of reproductive opportunities and the onset of mating) increases male lifetime reproductive success in a biologically relevant context (i.e. in competition against rival males over a series of different females; Figure 1). In contrast, we did not find evidence of net perception effects for intermediate sensory exposure treatments simulating a lag of 3d and 7d between the perception of reproductive opportunities and the onset of mating. Finally, and in line with previous evidence, we found that extended exposure to female cues prior to mating (i.e. 15d lag) led to net fitness costs in *D. melanogaster* males (5,7,22). Such fitness costs were due to a decrease in reproductive success (see also 22) but, contrarily to previous studies (5,7,22), we did not find that sexual perception reduced survival. This is likely due to the fact that our longest exposure (15 days) was considerably shorter than those in previous studies (see 5,7,22). To summarise, these results show that sexual perception is beneficial if males rapidly access mating after perceiving reproductive opportunities, and that sexual perception only leads to net fitness costs and accelerated ageing in males when the lag between perception of reproductive opportunities and the onset of mating is relatively long (i.e. 15 days long or more).

As population demography modulates the relative importance of reproductive timing (32–35), we complemented our analysis by calculating the rate-sensitive fitness consequences of male exposure to female cues. Our rationale was both to assess fitness effects in a range of demographic backgrounds and to examine the relative importance of early-versus late-life reproduction at modulating the aforementioned effects of sexual perception. While qualitatively similar across population dynamics, we found the potential for sexual perception to affect individual fitness to be more marked in decreasing populations, across all exposure lengths (Figure 2). Given that late-life reproduction is more important in decreasing populations, relative to stable or increasing populations (34, 35), this result highlights that sexual perception effects accumulate through life. Interestingly, post-hoc exploration of the data indicated that benefits linked to short-term (1d) perception were rapidly observable (as soon as over the 24 hours following the onset of mating) and persisted over the whole life of males (see SM for further details), implying that the physiological changes triggered by early-on perception of female cues were conserved in the long term. In other words, males do not seem to experience transient changes, but rather long-lasting responses that impact their life history. This is, in itself, a remarkable finding, and we suggest a priority for future studies should be to address the mechanism underlying this phenomenon. Understanding what specific fitness benefits (e.g. pre- vs. post-copulatory competition) are involved in these effects might offer valuable information about reproduction-survival trade-offs and the evolution of ageing.

The arising question is whether such male plasticity is likely to be adaptive for *D. melanogaster* males in nature, and available evidence strongly suggests it will. Virgin *D. melanogaster* females generally mate soon when presented to males, with average mating rates of one mating every 1 to 3 days in lab and wild populations (36–43). This implies that, despite strong male-male pre-copulatory competition and high variance in male mating success (44), most of the males that will ever reproduce are likely to start mating within their first two weeks of life. However, the aforementioned mating rates reflect situations of relatively high density that are more likely to represent maximum mating rates rather than to be indicative of average mating rates over space and time in the wild. While little is known about fine-grained population density dynamics in wild *D. melanogaster* (45), there is indirect evidence to suggest that density fluctuations are probably common in the field. This species’ ecology is closely linked to food sources (e.g. orchards) whose availability exhibits drastic spatiotemporal variation, which is inevitably bound to modulate local density. Accordingly, in the field *D. melanogaster* larvae maintain a stable polymorphism (largely driven by a single locus – the *for* gene–) with two foraging variants: a) rovers, characterized by long foraging trips and pupation away from the food source, are preferentially selected under high densities, while b) sitters, characterized by short foraging trips and pupation in the food source, are preferentially selected in low densities (46–47). Thus, there is suggestive evidence that natural populations of this species are subject to frequent fluctuations in local density. Under low densities and/or during phases where dispersal in search of food sources is likely to be common, finding (and mating with) females may be less frequent. This implies that average mating rates in the wild are almost certainly variable, as a consequence of fluctuations in local density. Under this context, male plastic responses such as those reported in this study are likely to be adaptive because they will allow males to engage the physiological machinery that allows them to maximize their competitive ability, but only in the presence of socio-sexual cues indicative of mating opportunities. Thus, a plastic response based on the presence of reliable socio-sexual cues seems to allow males to accrue the benefits of engaging in responses that condition them for competition over reproduction when in a high-density scenario, while avoiding the long-term costs that would ensue from unconditionally engaging such responses in a low-density environment.

A wealth of evidence has already accumulated to show that sensory cues, from nutritional to social cues, can profoundly modulate ageing in taxa as distant as flies and worms (21), but what do we know about their functional significance? It has been suggested that sensory modulation of ageing may result from a trade-off whereby organisms respond to ecologically relevant environmental cues to maximise early-life fitness at the expense of late-life fitness (21). However, we know little about the specific adaptive value of such plastic responses, and scarcely anything about what contexts we may expect these mechanisms to evolve in. In the case of ageing via sexual perception, an interesting facet of the costs involved in male plastic responses of this sort is that they are contingent on mating, so that costs are only paid if there is a relatively long lag between male perception of reproductive opportunities and the onset of mating (5,7,22). This has important implications for both the evolution of this strategy and its impact on related evolutionary processes. We suggest that male plastic behaviour similar to that reported here for *D. melanogaster* will arise frequently in nature in response to fluctuations in the availability of reproductive opportunities, whenever these correlate reliably with environmental cues (e.g. female odours). Implicit in this hypothesis are two predictions. First, that such changes must be, on average, beneficial to males facing imminent competition over reproduction, as strongly suggested by our results. Second, that male plastic responses should give rise to long-term physiological costs when males are unsuccessful, as demonstrated here and in previous studies reporting quite dramatic actuarial and reproductive costs in the case of *D. melanogaster* (5,7,22).

To date, ageing in response to cues of opposite sex has been reported in nematodes (*Caenorhabditis elegans*; 10) and flies (*D. melanogaster*; 5,7), suggesting that this phenomenon may be evolutionary conserved across the tree of life (3). Our simulations further show that male plastic responses to female cues will be favoured under a wide array of socio-sexual contexts. We found mating rate, as captured by the average β (onset of mating) in our simulations, to be the main determinant for the evolution of male plastic responses to reproductive cues. Consistently high mating-rates benefited a fixed “spendthrift” (i.e. always respond as if females were present) strategy (30-85% fitness advantage for mating rates of one every 1 to 4 days), while consistently low mating rates benefited a *“*thrifty” (i.e. always respond as if females were absent) strategy (38-56% fitness advantage for mating rates of one every 12 to 14 days; see SM for further details). In contrast, a plastic response would have higher fitness whenever average mating rates vary within an intermediate range of mating rates (Figure 4). Thus, we predict that in species where average mating rates are consistently high, such as promiscuous species with little fluctuation in density, a fixed spendthrift strategy will be favoured (Figure 4). This will be the case of species with very short lifespans, where long-term costs are likely to be negligible (mayflies are a good, albeit quite extreme, example). In contrast, we predict that a fixed thrifty strategy will be favoured in species with consistently low mating rates (Figure 4), such as iteroparous species with low density and/or prolonged reproductive seasons. Finally, male plastic responses are expected to evolve in species where mating rates fluctuate in accordance with changes in population density (Figure 4). Given that environmental stochasticity effects on population density are very frequent, we suggest that plastic responses to sexual perception, such as those reported here, might actually be a common strategy in nature; at least in promiscuous species with reproductive lifespans and demographic parameters within the range of *D. melanogaster* (modelled in this study), which are typical of many insects. Our model shows that under the latter condition, male plastic behaviour will be favoured almost irrespective of population demography (i.e. growth rate) and inter-individual variability in male mating rates. We thus suggest a priority for future studies should be to study the phenomenon of ageing via sexual perception across species with contrasting life histories.

**Figure 4:**
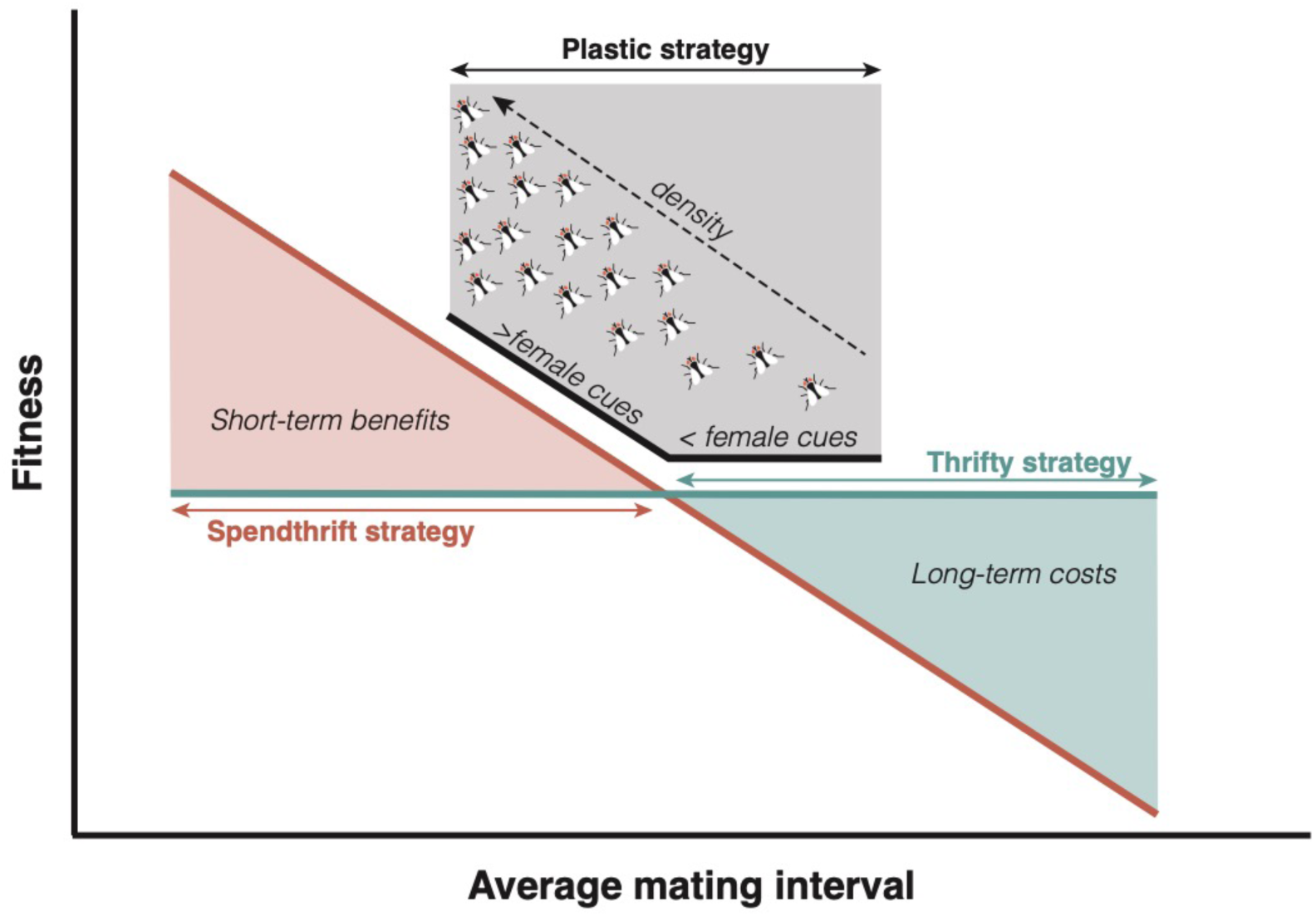
Alternative male strategies favoured depending on the range of variation in mating rates experienced by the population. Average mating rates that are consistently high will favour a fixed “spendthrift” strategy that taps on the short-term benefits of engaging a maximum male physiological response in preparation to competition for reproduction. In turn, average mating rates that are consistently low will favour a fixed “thrifty” strategy that avoids the long-term costs of engaging a maximum male physiological response in preparation to competition for reproduction. Finally, variation in mating rates within a low-to-moderate range (i.e. uncertainty as to whether males will be, on average, quick or slow to mate) will favour plastic male strategies whereby males engage (or not) maximum physiological responses depending on the presence of female cues (i.e. local mating patch density). This analysis assumes that local mating patch density correlates positively with average mating rates in the mating patch.

A second implication of the fact that male plasticity costs are contingent on mating is that this phenomenon can magnify sexual selection. The idea is that male responses to sexual perception can magnify sexual selection by further reducing the reproductive success of low-quality males. Briefly, if perception costs result from a decoupling between perceived and realised mating opportunities, it follows that low-condition males will tend to disproportionally pay such costs simply because they have lower mating success, and hence will take longer to mate (22, 23). In addition to this, our results suggest that males able to mate soon after perceiving female cues accrue a lifetime reproductive advantage over rival males. Given that high-quality males (i.e. good at intra-sexual competition) will tend to mate quicker than the average male, this means they are expected to disproportionally harvest perception benefits. As a consequence, male plastic responses to female cues are bound to increase the overall variability in male reproductive success (i.e. opportunity for selection), hence potentially magnify sexual selection beyond previously surmised. Our simulations show that this modulation can be biologically meaningful, with an average increase in the opportunity for selection estimated of between 15 and 27% for the whole range of average mating rates explored (Figure 4 & SM). Given that plastic male responses to female cues are expected to be favoured in promiscuous species, where sexual selection is already expected to be intense, this could lead to eco-evolutionary feedback that further magnifies sexual selection.

## MATERIALS AND METHODS

### Stocks and maintenance

Unless when stated otherwise, all flies used in this experiment were laboratory wild type (*wt*) Dahomey *D. melanogaster*. We used homozygous recessive *spa* mutants (*sparkling poliert*) as competing males and reproducing females in order to assess paternity of focal *wt* individuals. Homozygous *spa* flies have a distinguishable rough-eye phenotype that allowed us to distinguish all offspring from our focal males, and have the added advantage of being slightly worse competitors than *wt* males, which ensured focal males would eventually mate. Stock populations are maintained outbred, with overlapping generations, at 25°C on a 12h light/12h dark cycle fed with standard food (solidified aqueous mix containing 60g.L^−1^ corn flour, 50g.L^−1^ white sugar, 40g.L^−1^ fresh baker’s yeast, 10g.L^−1^ soy flour, 10g.L^−1^ industrial agar, 3g.L^−1^ Methyl 4-hydroxybenzoate (nipagin), 10mL.L^−1^ 96 % EtOH, 5mL.L^−1^ 99% propionic acid). We obtained all flies by collecting eggs on yeasted grape juice agar plates (FlyStuff grape agar premix, Genesee Scientific) from stock populations. We reared all flies used in this experiment at a controlled density of ca. 200 individuals per 250mL bottle filled with ca. 75mL of food, and isolated them by sex within 6 hours of emergence (i.e. as virgins) at standard densities of 15 females and 20 males per vial, using ice anaesthesia.

### Experimental design

We individually exposed *wt* males to either donor females (treatment) or not (control) following the method as described in García-Roa *et al.* (22). Briefly, we connected two vials containing food to each other and placed a mesh partition between them. Males were then singly isolated on one side of the mesh, while the other contained either: a) three *wt* females (i.e. treated males) or b) no females (i.e. control). This mounting allowed treated males to be exposed to female odours while ensuring they would not mate. To explore whether sensory perception effects are contingent on the time of exposure to female cues, we exposed experimental males to sensory stimuli (i.e. female cues or control) for one day, three days, seven days or 15 days; n = 60 per treatment combination (i.e. 2 treatments * 4 exposure times * 60 biological replicates, for a total of 480 focal flies). This replicate number was determined based on the effect size obtained in a prior experiment (22) exploring the effects of sexual perception on reproductive success.

Immediately following sensory exposure treatments, we transferred focal males into vials in which they competed over a *spa* female against two *spa* males for the rest of their lifetime (all four flies were virgin at the beginning of assays). During this time, we checked for survival daily and replaced competitors and females that died before our focal male by virgin individuals of the same age. We also swapped competitors and females for younger virgin individuals (less than 12 days old) every 10-11 days to simulate a natural situation, where males have to compete for different females over their lifetime. We transferred flies into new vials every 3-4 days, and incubated vacant vials for 12 days (time needed to ensure all viable flies have emerged, given that average generation time is 8 to 9 days), before freezing them at −20°C for later counting of *wt* and *spa* F1 offspring. Treatment allocation was masked during counting. For transfer of flies from one vial to another one, we used gentle CO_2_ anaesthesia. All steps of the experiments were performed at 25°C.

### Statistical significance analyses

We used Cox proportional hazards (48) for survival and post-hoc reproductive lifespan analyses, checking assumptions using statistical and graphical diagnostics based on Schoenfeld residuals (49). We fitted a general linear model (LM) in order to analyse relative reproductive success (using *wt/*total number of F1 – #F1–).To analyse reproductive ageing, we first calculated the average reproductive success of each focal male over each one of the first three females it competed for (only for individuals surviving this period of time). Then, we calculated the net decrease in male relative reproductive success between the first and the second female (Δ♀_1_♀_2_), as well as between the first and the third female (Δ♀_1_♀_3_). We fitted general linear models to Δ♀_1_♀_2_ and Δ♀_1_♀_3_.

All these models incorporated treatment (categorical variable, two levels: sensory sexual perception and control), exposure time (continuous variable) and the interaction between treatment and exposure as fixed factors. We checked model assumptions (residuals normality, homoscedasticity, homogeneity of variances, absence of outliers and influential points, absence of autocorrelation of factors) using the “performance” package (50). We assessed model term significance with α=0.05. Models were run computing type III ANOVA using the “car” package (51), in R studio 1.1.456. No clear outlier was detected in the dataset, and no datapoint was excluded of statistical analyses. All significant reported p-values remain so after correcting for inflation of type I error rate due to multiple testing (using the Benjamini-Hochberg procedure for a false discovery rate of 0.05).

In order to place lifetime reproductive success effects in a demographic context, we additionally estimated individual rate-sensitive fitness estimates of treated versus control individuals across exposures (1, 3, 7 and 15 days) for different population growth rates (r = −0.1, - 0.05, 0, 0.05 and 0.1; 32,33,52). For the latter analysis, we included results from García-Roa *et al.* (22) as a historical treatment/control, as this study used the same experimental design as well as the same lab population of flies, but with a sensory perception exposure time of 21 days.

### Modelling and simulations

Following Tatar & Promislow (53) we define fitness as the integration over age from x = 0 to ω (the end of reproductive ages) of

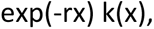

where r is the intrinsic growth rate, and k(x) is the so called reproductive function (l(x) m(x); i.e. survival times fertility at age x; e.g. 33). Additionally, x = 0 is the age at maturity, so that developmental times are not considered to make a difference when fitness is compared between scenarios or individuals. Our computer simulation model is informed by our experimental design, but making the biological mechanisms as explicit as possible and using directly interpretable factors that act on k(x). In this way, the values of model parameters can be obtained from the estimates from experimental data.

Therefore, in order to build our computer simulation model, we performed data analysis to describe and extract model components. In that analysis, k(x) was analysed on an individual basis; i.e., k_i_(x), where i identifies the i^th^ experimental individual. The realized k_i_(x) is a count; i.e., the number of offspring at age x (i.e. from x to x + Δ_x_x; the subscript stresses that the experimental time lag between consecutive observations is not constant) of the individual i^th^. Commonly in Mixed Effects Models the expectation of the log-count is modelled additively; in our case

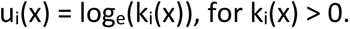

As u_i_(x) may be a rather complex function, we used Generalized Additive Models (GAMs). We assumed a negative binomial (NB) error distribution and applied GAM separately for each experimental condition (four treatments and four controls). In addition to a structural relationship between u_i_(x) and x (a more or less elevated smooth function to be found), our GAM analysis assumed two random components (random slope and random intercept; both following uncorrelated Gaussian distributions) to account for among-individual variation in the experiments. We used packages mgcv (54) and nlme (55). GAMs found smooth functions (expected u_i_(x) vs. x) with similar shapes (approximately, an asymmetric a 90-degree rotated sigmoid curve) for the eight experimental conditions (see SM for further details. The shape was compressed for the life window with potential access to females; that is, between the onset of access to females (B; 1, 3, 7 and 15 days) and the end of the reproductive life. The rotated sigmoid functions were more or less elevated depending on the experimental condition, and we found a clear pattern whereby elevation decreases with the female access onset. Using least squares, we fitted lines (treatment and control separately) to the relationship between the elevation and the female access onset (E vs. B). Our GAMs did not find random intercepts to be significant but did find a significant variation in random slopes.

After this analysis, for our computer model the i^th^ individual is assumed to have the expected exp(u_i_(x)) (i.e. randomness from NB distribution was neglected). Our computer model uses the functional shape showed in SM with the elevation describes by SM). However, in order to compute elevation of u_i_(x) now the mating onset is β (not B, fixed experimentally) and is a log-normal variable with values assigned randomly to males. Additionally, a male-dependent Gaussian random slope is added to get u_i_(x). We envisage the random slope as being due to demographic stochasticity, and assumed it to be independent on the timing of the mating onset. Simulations based in 10000 were replicated twice, and performed using Wolfram Mathematica code (release 10; Wolfram Research, Inc.). Additional technical details and parameter values are provided in the SM.

## ACKNOWLEDGEMENTS

We thank Claudia Londoño, Alejandro Hita and Mariona Vinyeta for help with experimental set up, data collection and data entering, respectively.

## DATA AVAILABILITY

Dataset uploaded to Dryad repository url: https://doi.org/10.5061/dryad.sn02v6x3w

## AUTHORS’ CONTRIBUTIONS

QC, RG-R and PC conceived and designed the study. QC conducted the experiment with help from RG-R and PC. QC analysed the data. MS conceived the theoretical framework for the mathematical models with help from PC. MS built the mathematical models and ran the simulations. QC and PC wrote the manuscript with input from MS and RG-R. All authors approved the final version of this manuscript. Authors declare no competing interests.

## Supplementary material

**Supplementary 1.** Table displaying F-statistic and *P*-value of the exposure*treatment interaction for population growth-rates ranging from −0.1 to 0.1. Only exposure times ranging from 1 to 15 days were used in this analysis. The exposure*treatment interaction is clearer at negative growth-rate than it is at positive growth-rate. This demonstrate that late-life reproduction is of crucial importance in the effects observed.

|  | Population growth-rate (*r*) |  |  |  |  |
| --- | --- | --- | --- | --- | --- |
|  | *r* = - 0.1 | *r* = - 0.05 | *r* = 0 | *r* = 0.05 | *r* = 0.1 |
| *F*<sub>(3, 405)</sub> | 2.545 | 2.462 | 2.287 | 2.087 | 1.897 |
| *P*-value | 0.056 | 0.062 | 0.078 | 0.102 | 0.130 |

**Supplementary 2.**
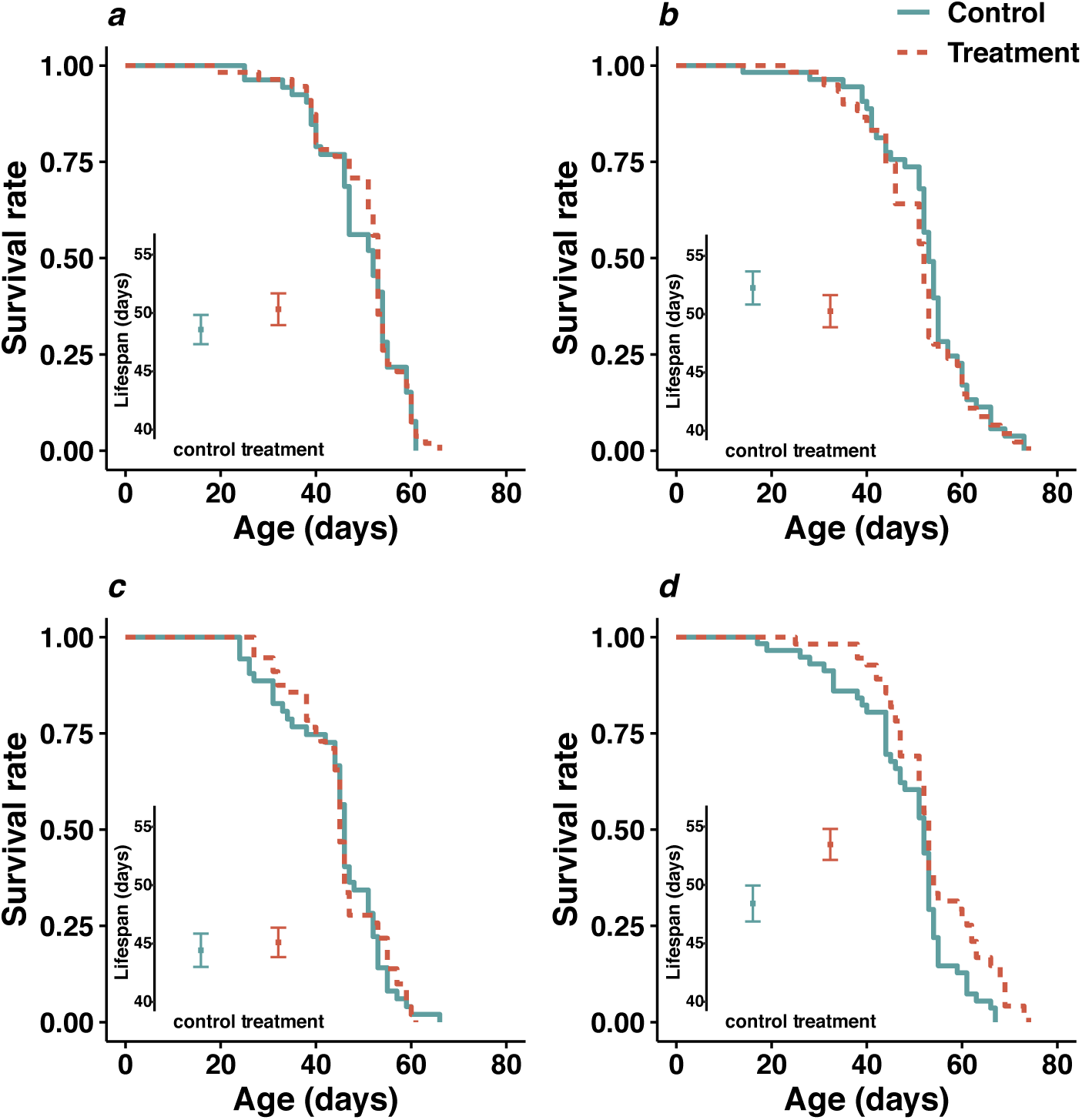
Survival rate as a function of age and average lifespan of treated and control individuals following 1 day (**a**), 3 days (**b**), 7 days (**c**) and 15 days (**d**) exposure. The orange dashed line represents the survival of treated males and the green continuous line represent the survival of control males. Green and orange squares depict the average lifespan of treated and control males, respectively, and vertical bars represent the standard errors around the average lifespan.

**Supplementary 3.**
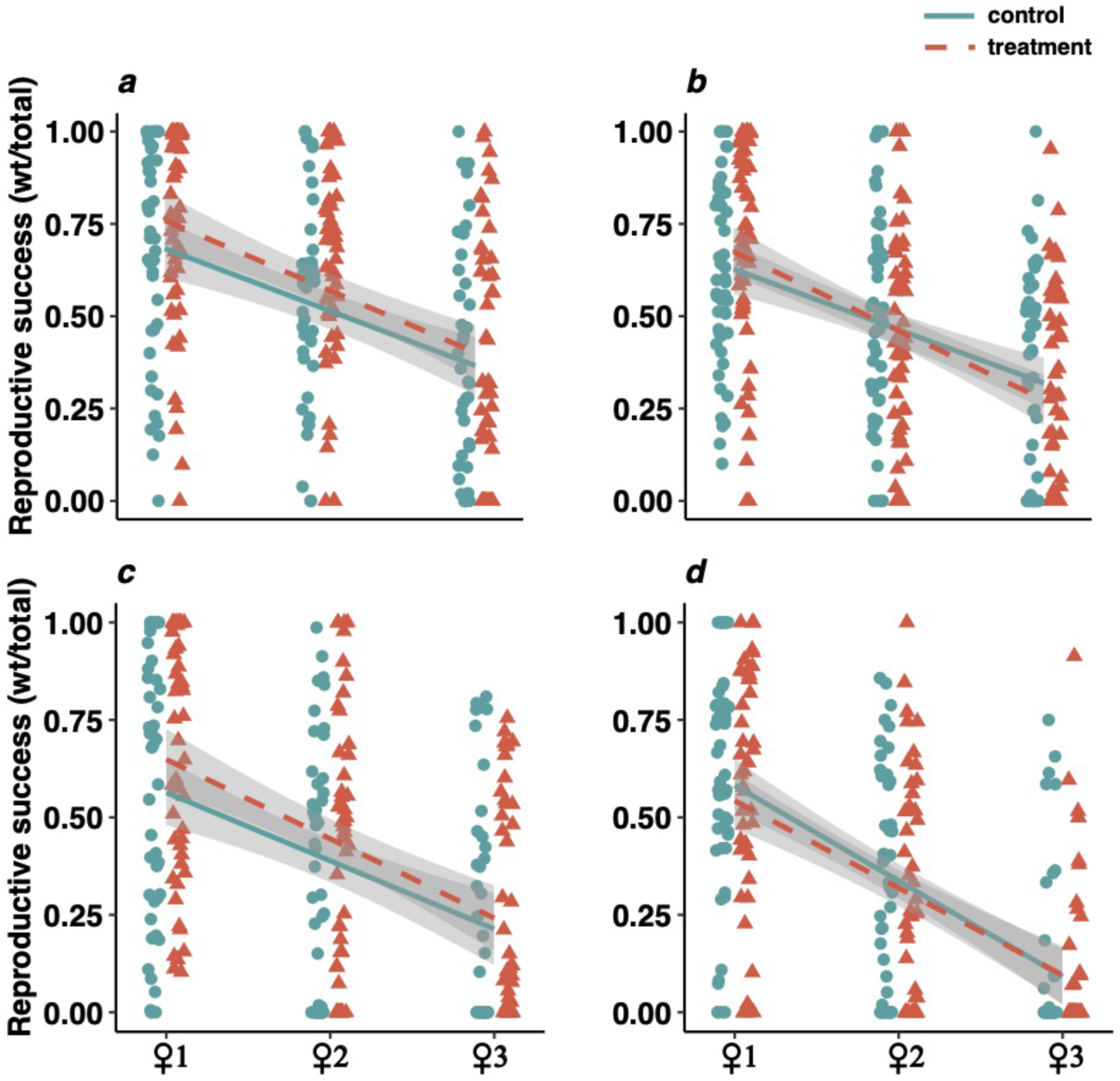
Relative reproductive success (percentage of offspring fathered by the focal male) of treated and control males, following 1 day (**a**), 3 days (**b**), 7 days (**c**) and 15 days (**d**) exposure, for the first three females they were competing over. Each orange triangle represents an observation of a treated male (with sexual perception) relative reproductive success over a female (12 days), while green circles are for control males. The linear regression lines represent the rate of decrease in relative reproductive success with the orange dashed line corresponding to treated males (and the green continuous line is the regression corresponding to control males, over the first three females they were competing over. Greyed areas around the regression lines represent 95% confidence intervals.

**Supplementary 4.**
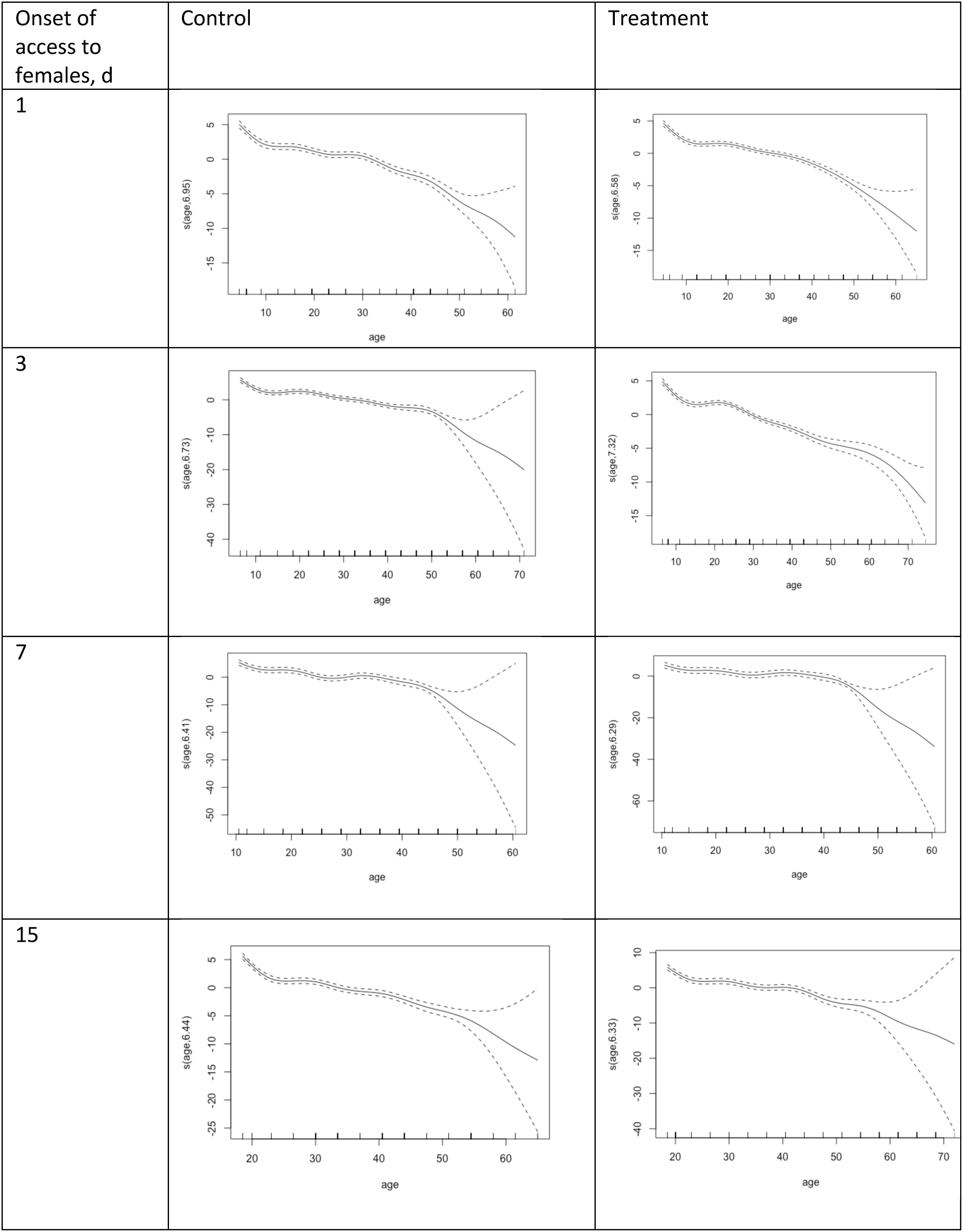
Smooth functions found by GAM and the shape of log-reproductive function for female sensing males (treatment) and control males, across exposure times (1, 3, 7, 15 days)

**Supplementary 5.**
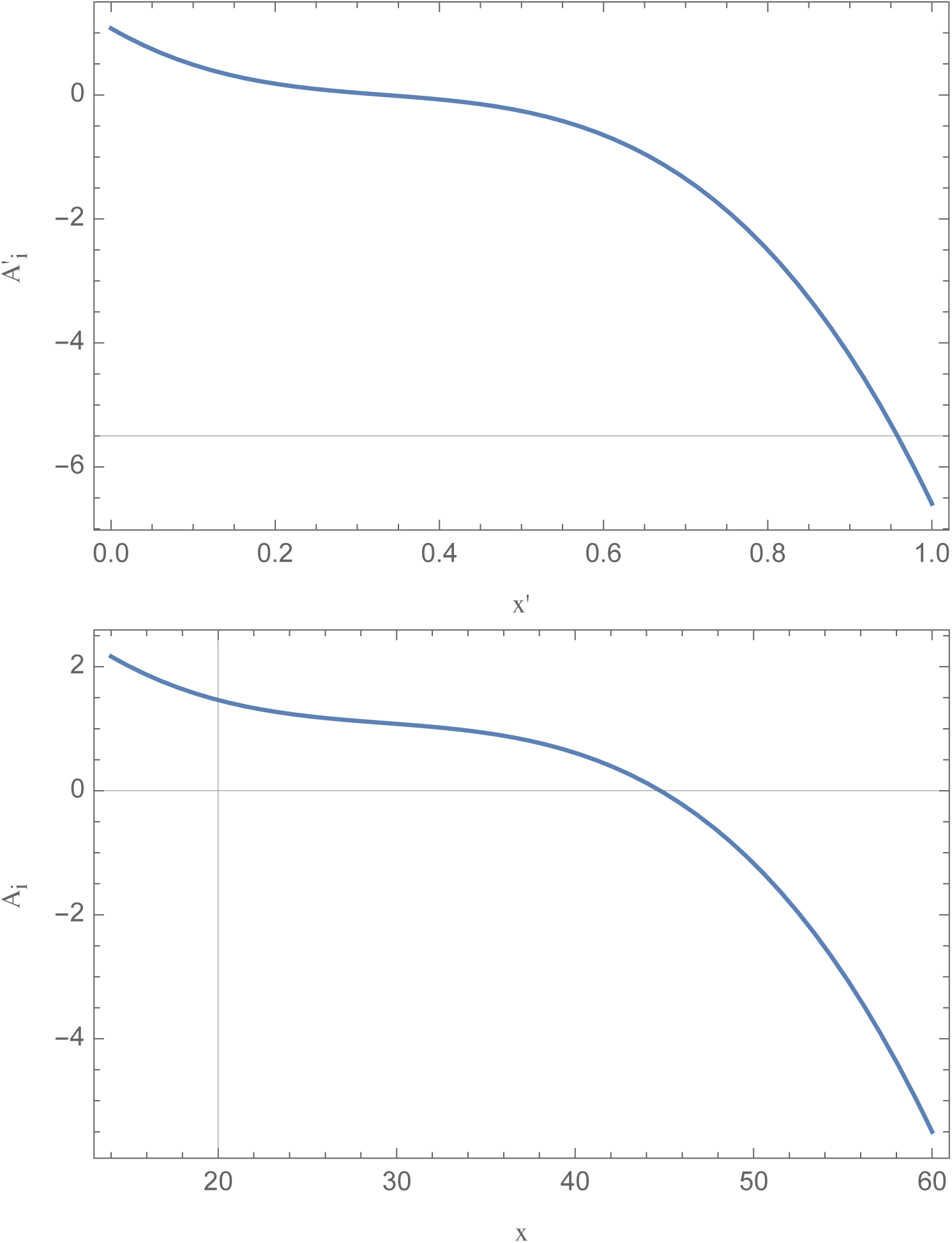
Log-reproductive function. The showed shape is assumed in the computer model for the structural component of the log-reproductive function. Upper panel, core function (before rescaling; b1 = −1; b3 = −20; c = 1/3. Lower panel, rescaled function for the case β_i_ = 14 and ω = 60; note that the average value for the function in the realized range is equal to zero. The structural part of the log-reproductive function is completed by adding a constant value, the elevation (E). In order to get the log-reproductive function of a male, a linear component with random slope is added.

**Supplementary 6.**
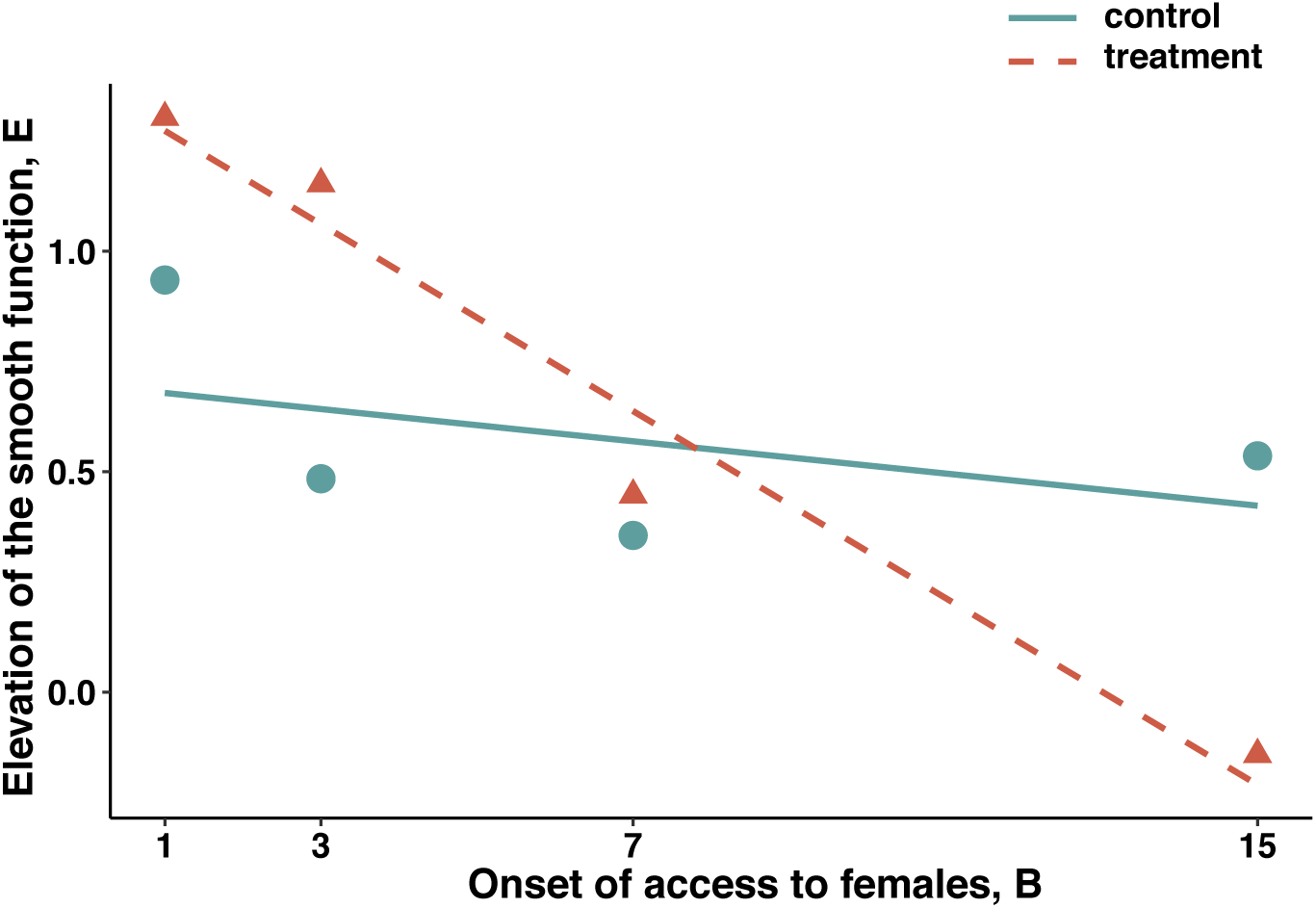
Relationship between the elevation (E) of the smooth function found after GAM and the experimentally controlled onset of access to females (B, age in days). Lines fitted using LS regression. Treatment: E = −0.11 B + 1.379, R^2^ = 0.96, P= 0.019. Control: E = −0.02 B + 0.699, R^2^ = 0.21, P=0.546. These lines were used in our simulations to model the “spendthrift” and “thrifty” strategies, respectively, but substituting B (access to females; experimentally fixed, common to any male) for β (mating onset; randomly chosen from a lognormal distribution for each simulated individual).

**Supplementary 7.**
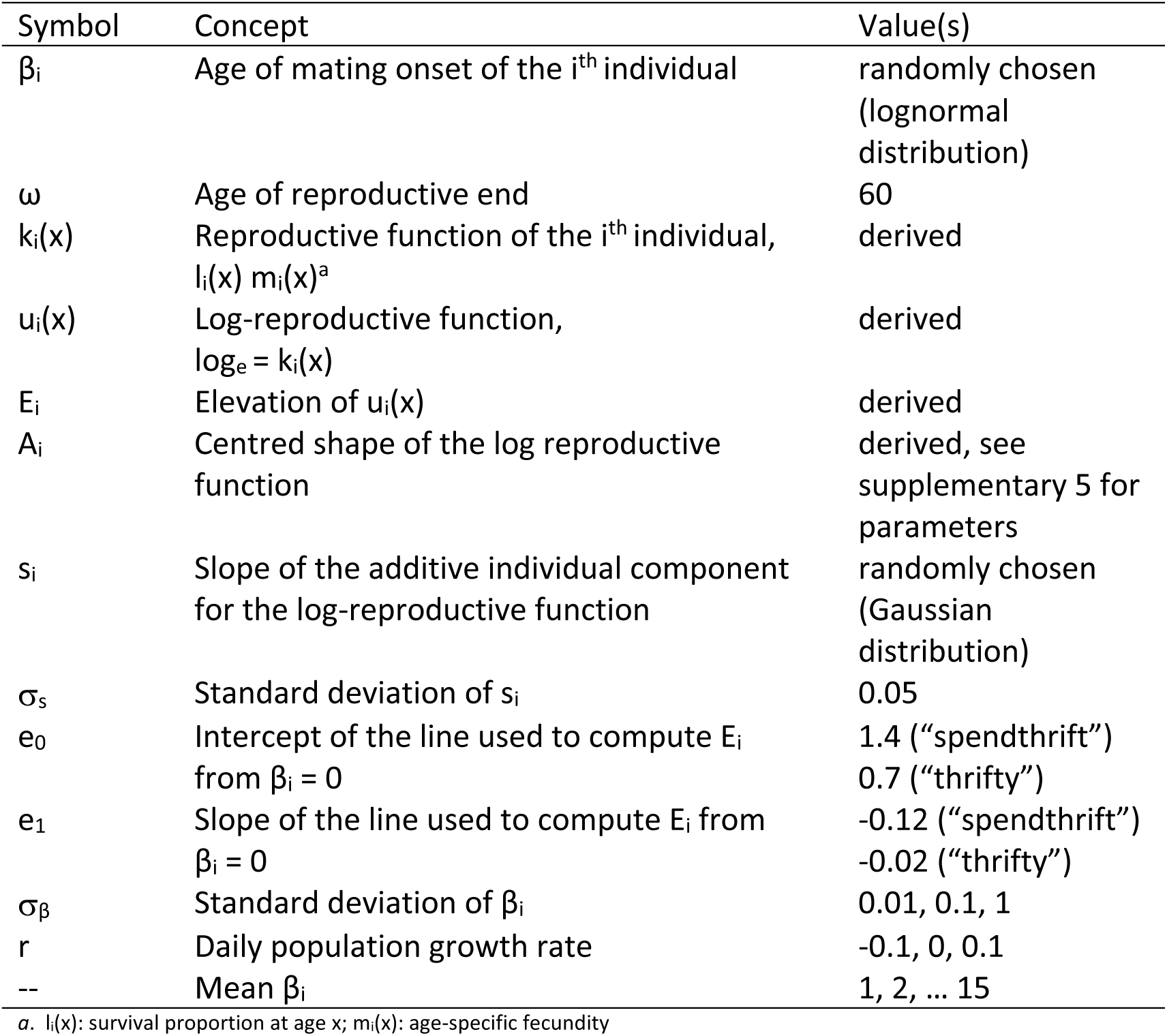
Computer model components and values

| Symbol | Concept | Value(s) |
| --- | --- | --- |
| $\beta_i$ | Age of mating onset of the $i^{\text{th}}$ individual | randomly chosen<br>(lognormal<br>distribution) |
| $\omega$ | Age of reproductive end | 60 |
| $k_i(x)$ | Reproductive function of the $i^{\text{th}}$ individual,<br>$l_i(x) m_i(x)^a$ | derived |
| $u_i(x)$ | Log-reproductive function,<br>$\log_e = k_i(x)$ | derived |
| $E_i$ | Elevation of $u_i(x)$ | derived |
| $A_i$ | Centred shape of the log reproductive<br>function | derived, see<br>supplementary 5 for<br>parameters |
| $s_i$ | Slope of the additive individual component<br>for the log-reproductive function | randomly chosen<br>(Gaussian<br>distribution) |
| $\sigma_s$ | Standard deviation of $s_i$ | 0.05 |
| $e_0$ | Intercept of the line used to compute $E_i$<br>from $\beta_i = 0$ | 1.4 ("spendthrift")<br>0.7 ("thrifty") |
| $e_1$ | Slope of the line used to compute $E_i$ from<br>$\beta_i = 0$ | -0.12 ("spendthrift")<br>-0.02 ("thrifty") |
| $\sigma_\beta$ | Standard deviation of $\beta_i$ | 0.01, 0.1, 1 |
| $r$ | Daily population growth rate | -0.1, 0, 0.1 |
| -- | Mean $\beta_i$ | 1, 2, ... 15 |

a. $l_i(x)$ : survival proportion at age $x$ ; $m_i(x)$ : age-specific fecundity

**Supplementary 8.**
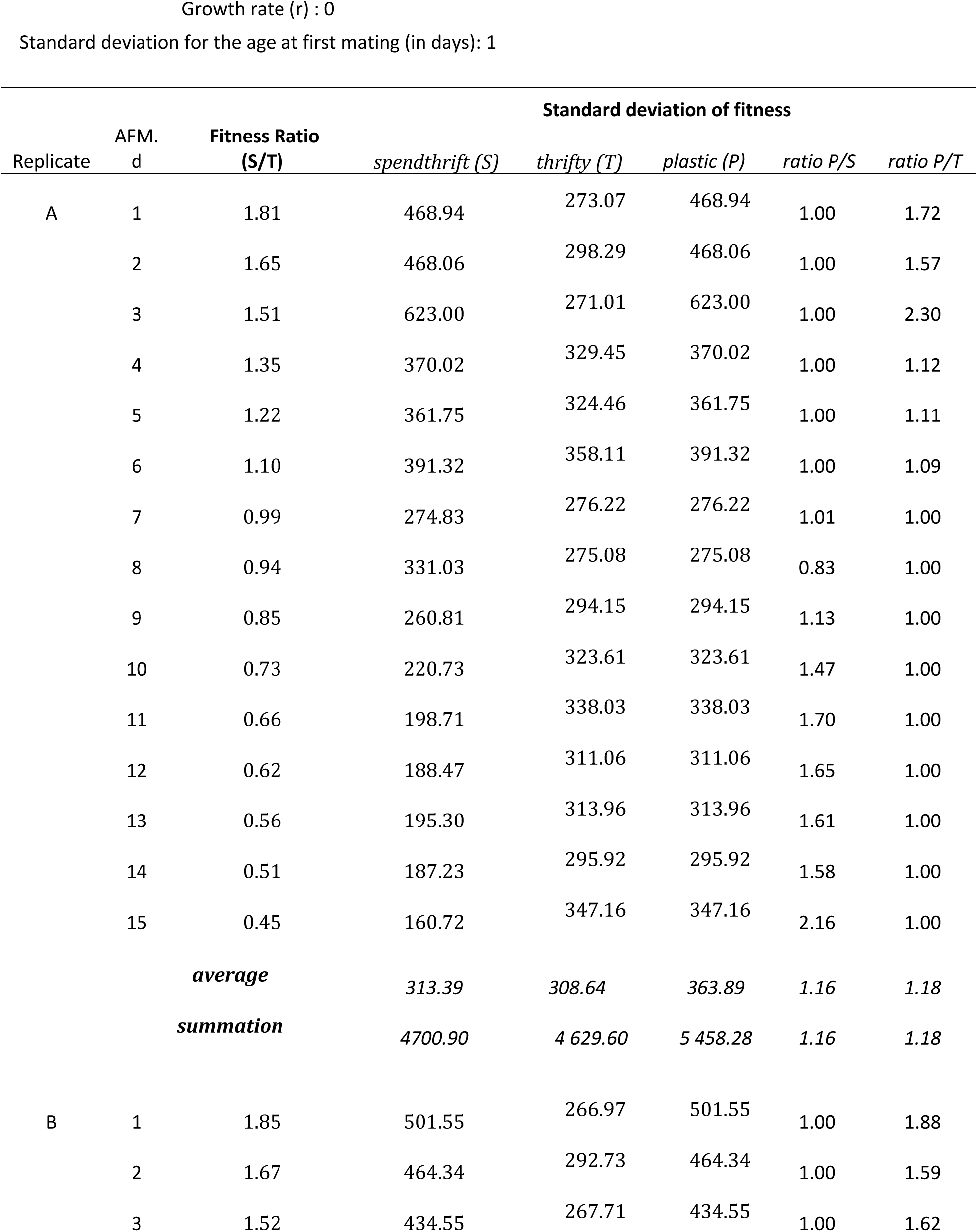
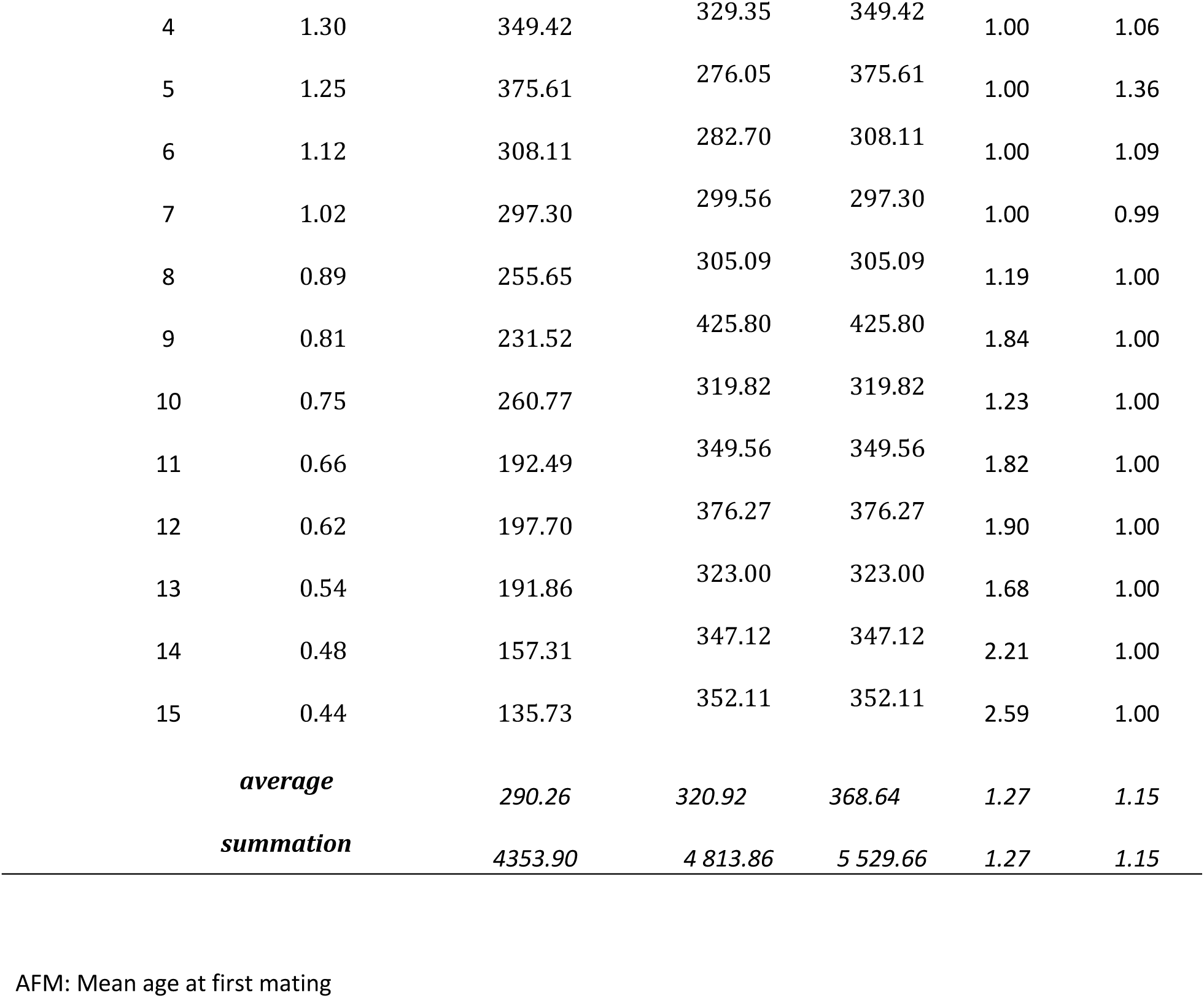
Simulation results for the two replicates that assumed r (daily growth rate) = 0 and standard deviation of mating onset (in days) = 1. Each replicate is based on n= 10000 males.

**Supplementary 9.**
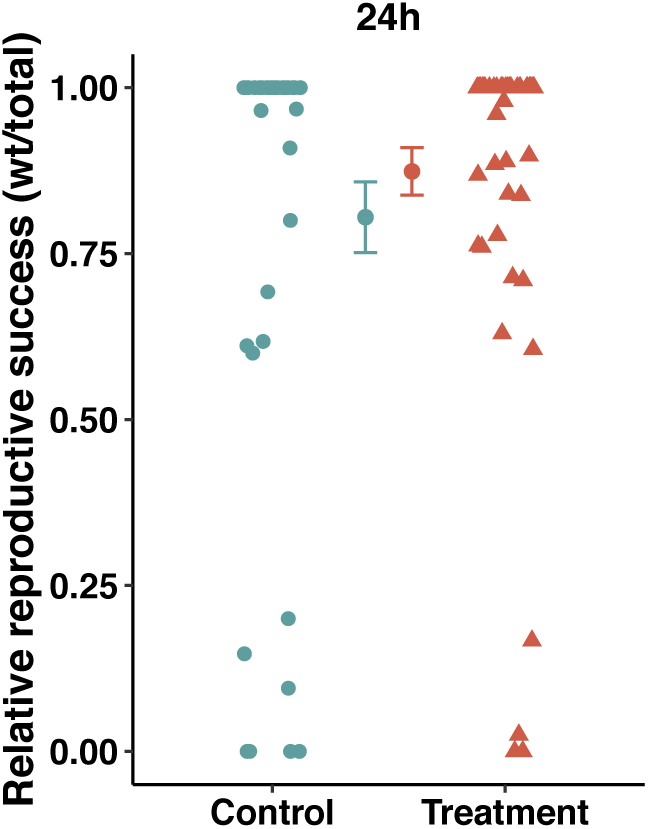
Mean relative reproductive success of control (green circles) vs treatment (orange triangles) males over the first 24 hours of competition for reproduction against two rival males for access to female, with the onset of access to female manipulated to 1 day. Vertical bars represent the standard error around the mean

**Supplementary 10.**
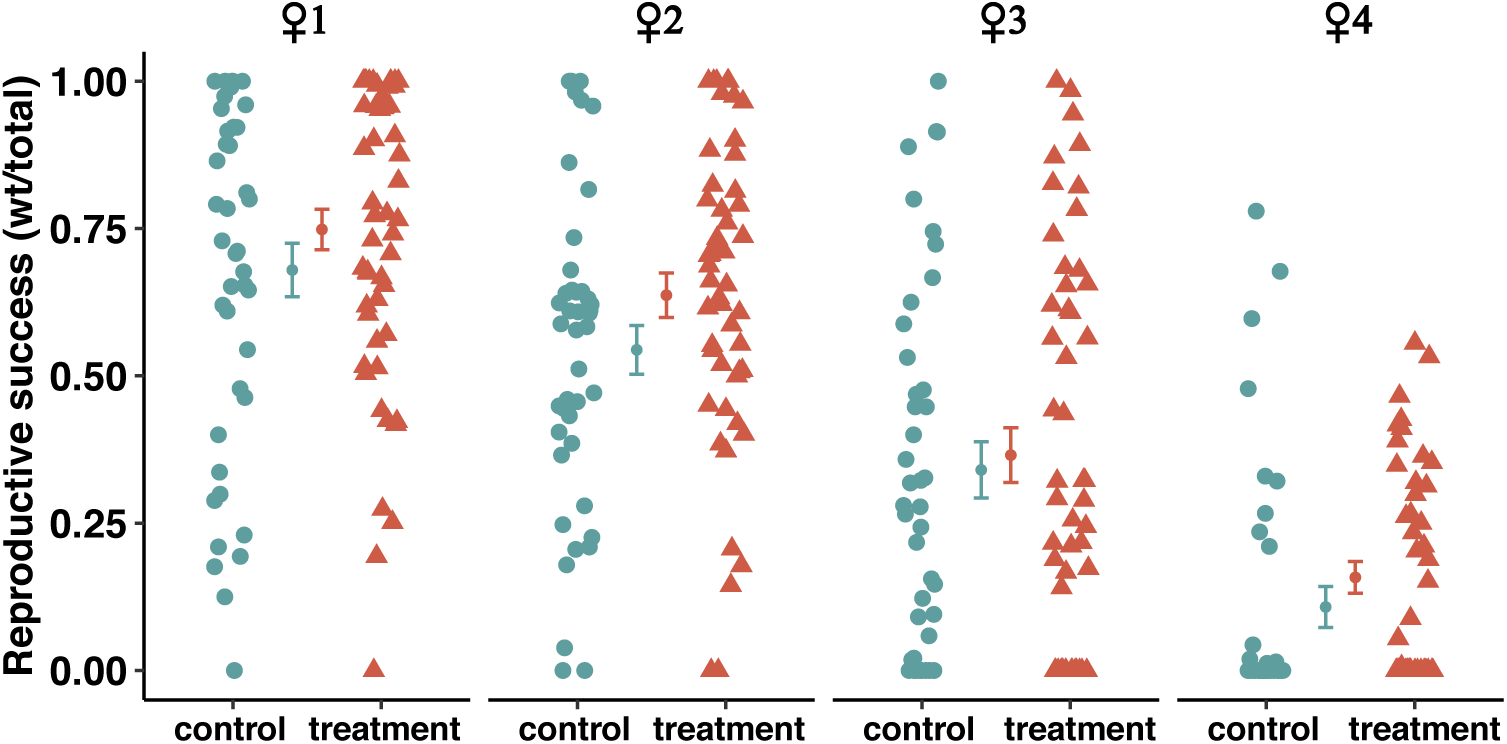
Mean relative reproductive success of control (green circles) vs treatment (orange triangles) males over the first, second, third and fourth female males were presented to, with the onset of male access to female manipulated to 1 day. Only males surviving to compete over the corresponding female are represented (i.e. zero values show males accomplishing 0% reproductive success). Vertical bars represent the standard error around the mean.

**Supplementary 11:** additional Material and Method with regards to computer model and simulations

1. k_i_(x) = 0, for x < β_i_, and k_i_(x) = exp(u_i_) for x ≥ β_i_, where β_i_ is the age at first mate of the i^th^ individual.
2. β_i_ is assumed to be a random variable with

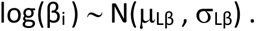 Note that the mean of β_i_ increases with σ_Lβ_, and conveniently, if μ_Lβ_ = σ_Lβ_ = 0, β_i_ = 1 for any individual. We computed μ_Lβ_ and σ_Lβ_ from the parameters at the arithmetic scale, μ_β_ and σ_β_.
3. For x ≥ β_i_, u_i_(x) is the summation of three components: elevation (E_i_), structural deviation [A_i_(x)] due to reproductive age, and individual (random) deviation within the effective reproductive period (s_i_). We used subscripts for E and A because they are assumed to be β_i_-dependent (see below).
4. We modelled E_i_ as a linear function of β_i_ according to the relationships depicted in supplementary 5, that is,

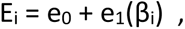

where e_0_ and e_1_ are the parameters of the linear relationships. Two scenarios are assumed: females perception is conditioning the male response (“spendthrift” strategy, henceforth *S* strategy) or not (“thrifty” strategy, henceforth *T* strategy), and accordingly two pairs {e_0_, e_1_} were be used.
5. A_i_(x) is a rotated sigmoid function of x for x ∈ [β_i_, ω[. We chose a simple (cubic) function based on the inspection of the smooth functions found by GAM (Supplementary 4). Our function is a linear modification of

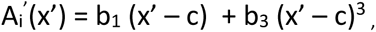

(core function) where b_1_ and b_3_ are the polynomial (negative) coefficients and x’ ranges from 0 to 1. When x’ = c the function has its maximum slope (= b_1_). The equation is rescaled from x’ to x, which ranges from β_i_ to ω. By normalizing A_i_^’^(x) so that it is centred to zero for x ∈ [β_i_, ω [, we obtain A_i_(x). We used parameter values consistent with our empirical observations, giving priority to early ages, since late ones are expected to have low effect on fitness.
6. s_i_ is modeled as a random Gaussian slope for the relationship between u_i_ and x, which accounts for individual deviations from A_i_(x). That is,

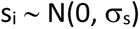 In summary, our model is k_i_(x) = 0, for x ∈ [0, β_i_[, log[k_i_(x)] = e_0_ + e_1_(β_i_) + b_1_ ([x – β_i_] / [ω - β_i_] – c) + b_3_ [x – β_i_] / [ω - β_i_] – c)^3^ - 1/[ω - β_i_] ∫_βi_^ω^ b_1_ ([y – β_i_] / [ω - β_i_] – c) + b_3_ (y – β_i_] / [ω - β_i_] – c)^3^ dy + s_i_ x, x ∈ [β_i_, ω [(y: age); s_i_ ∼ N(0, σ_s_); log(β_i_) ∼ N(μ_Lβ_, σ_Lβ_). See supplementary 4 for parameter values used in computer simulations.

**Supplementary 12:**
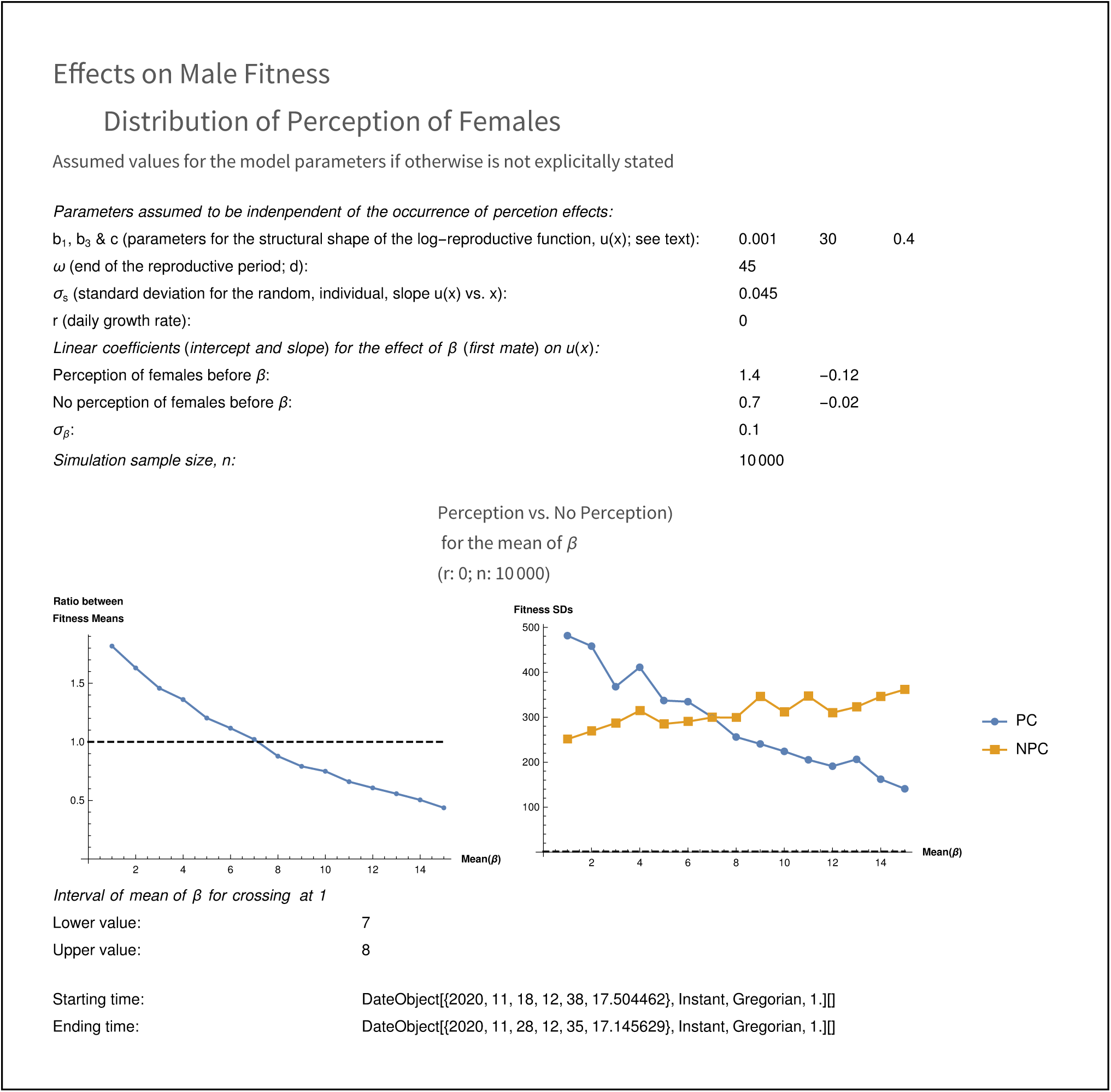
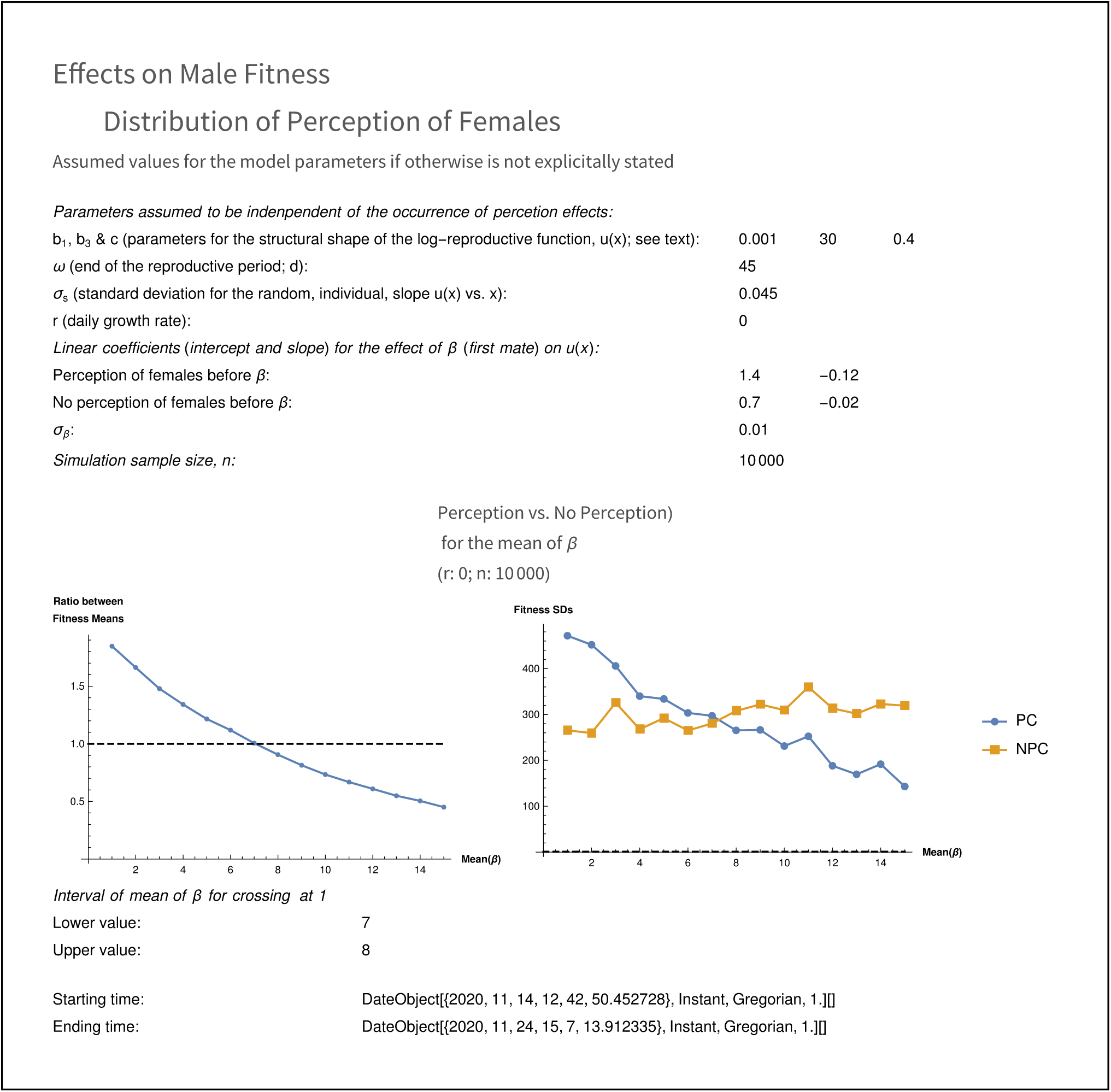
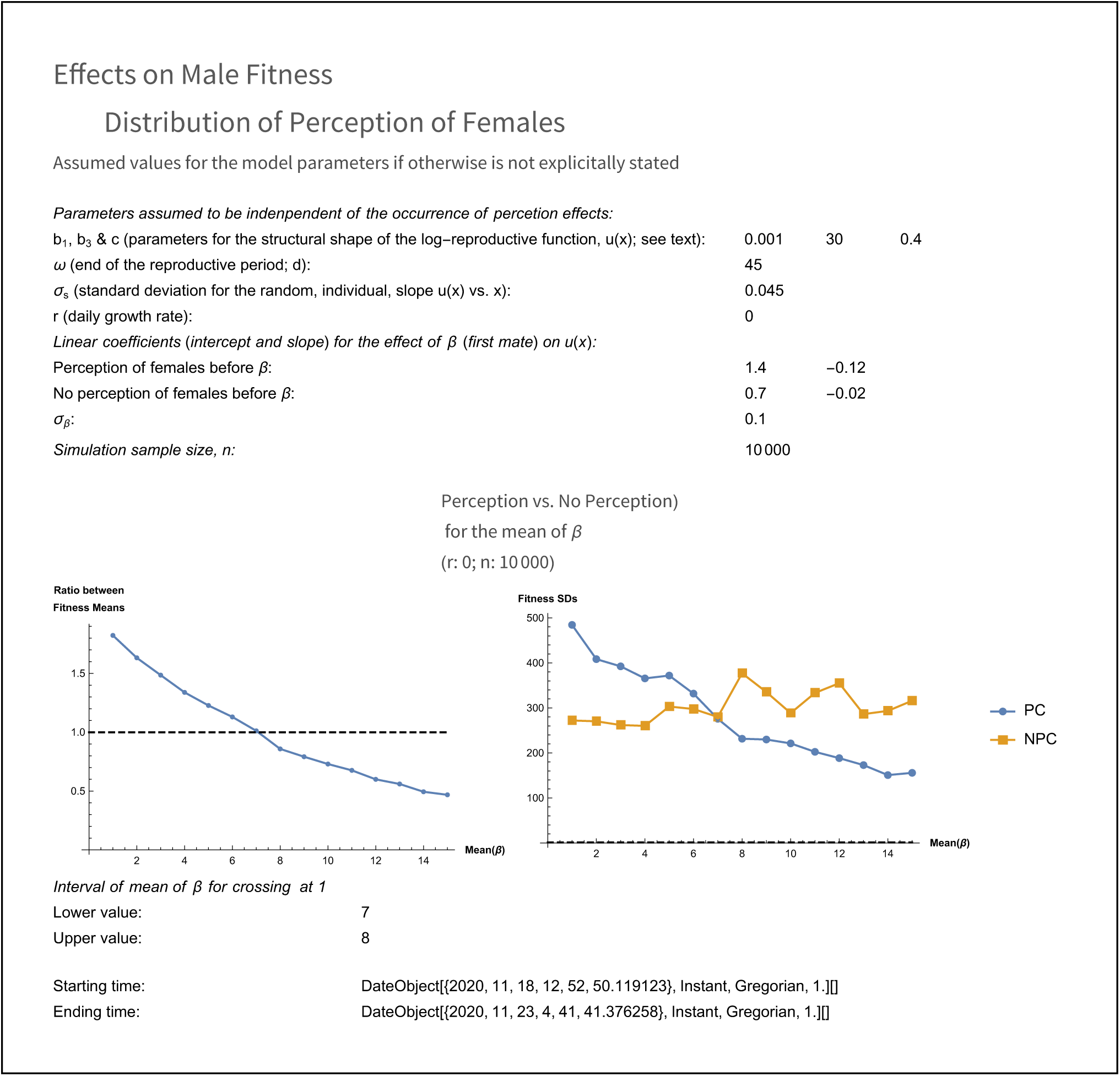
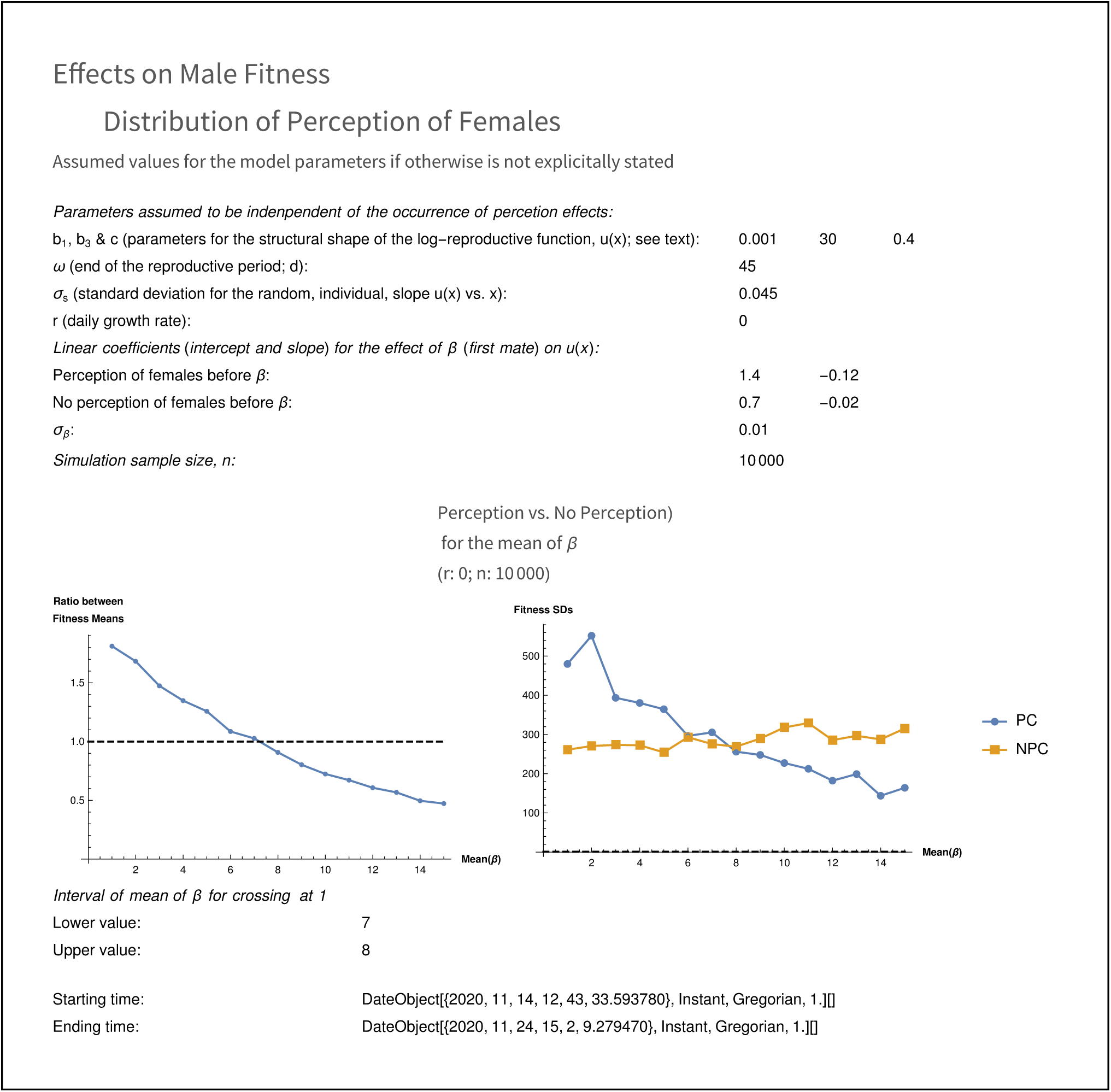
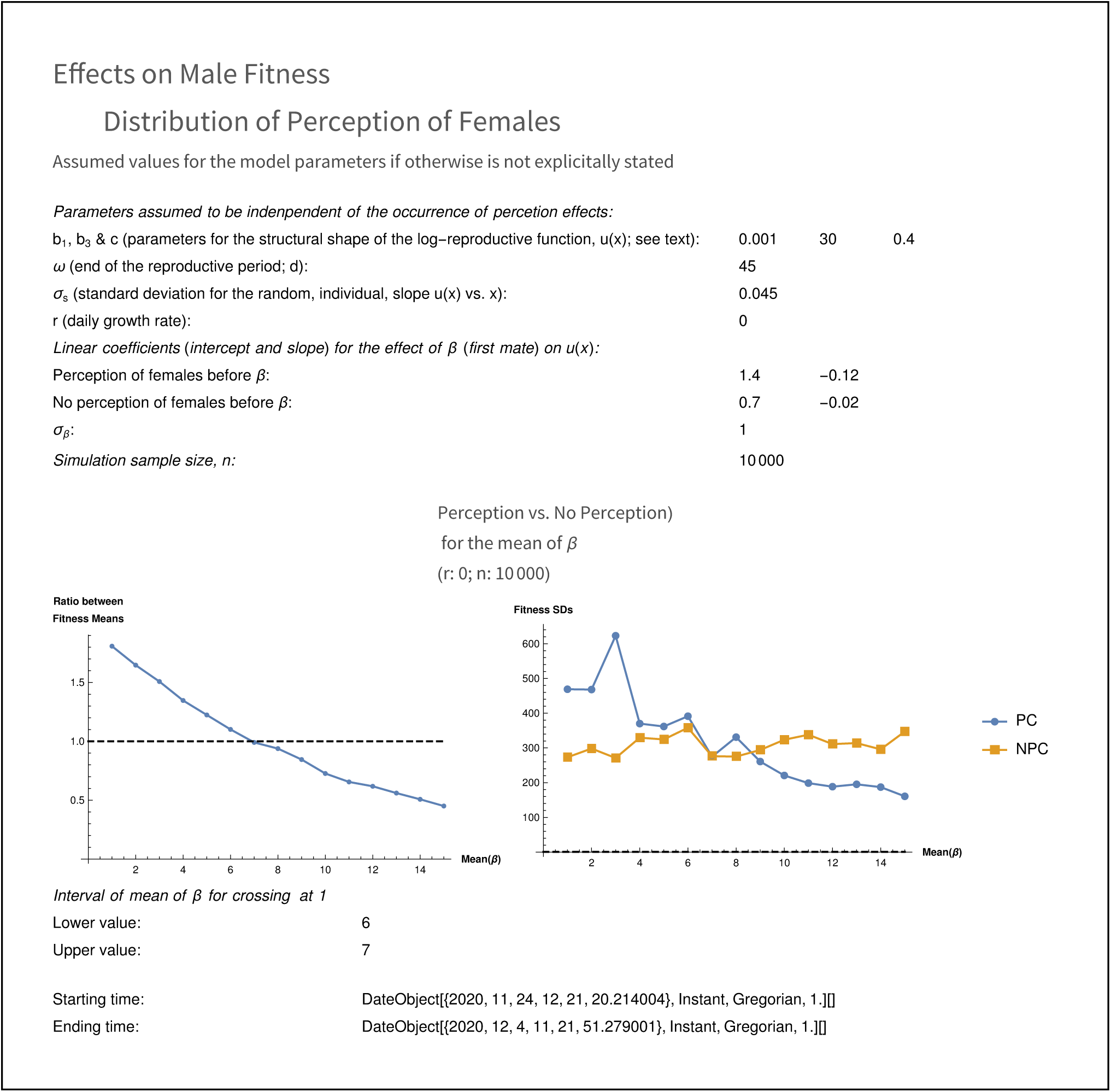
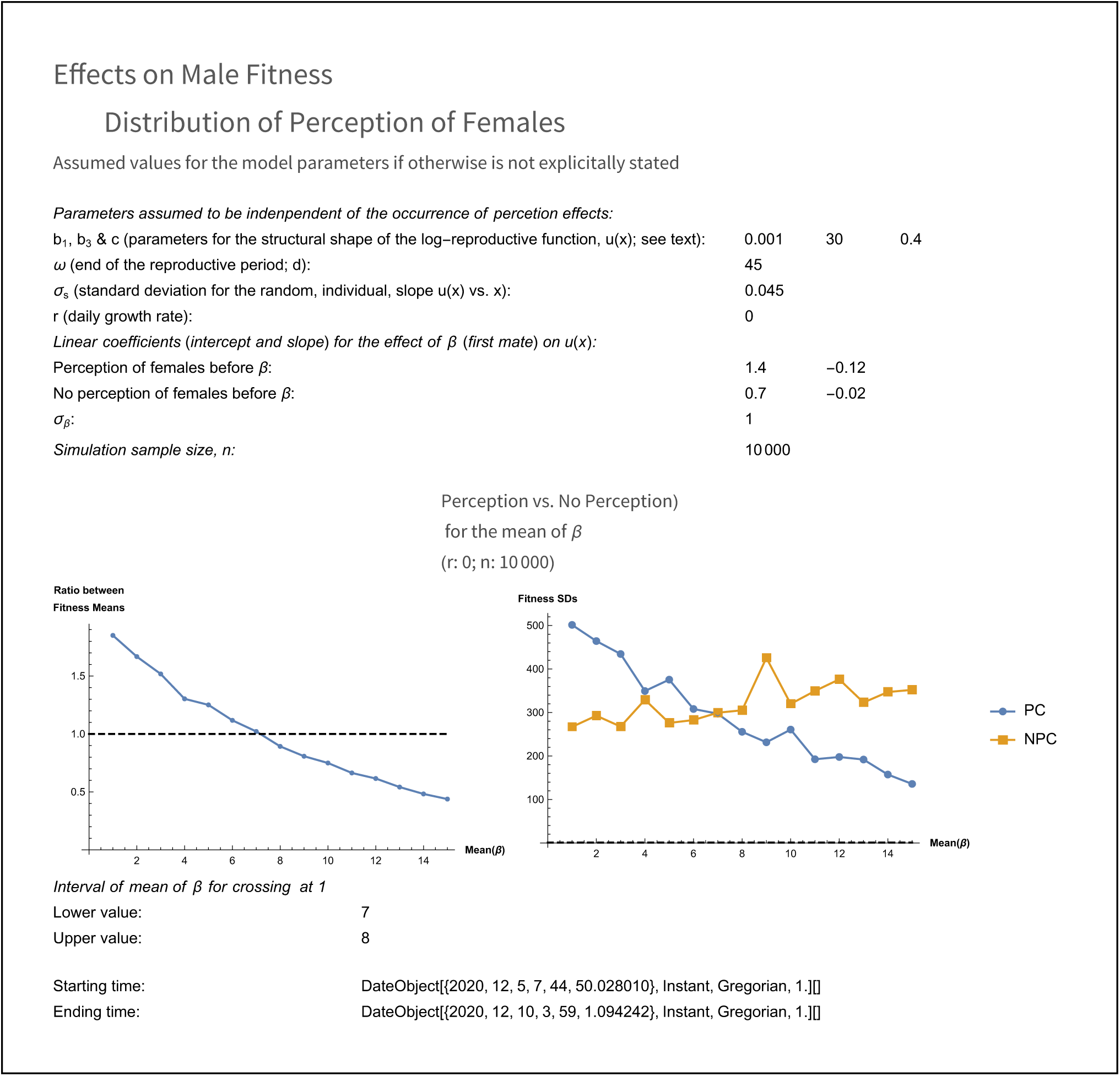
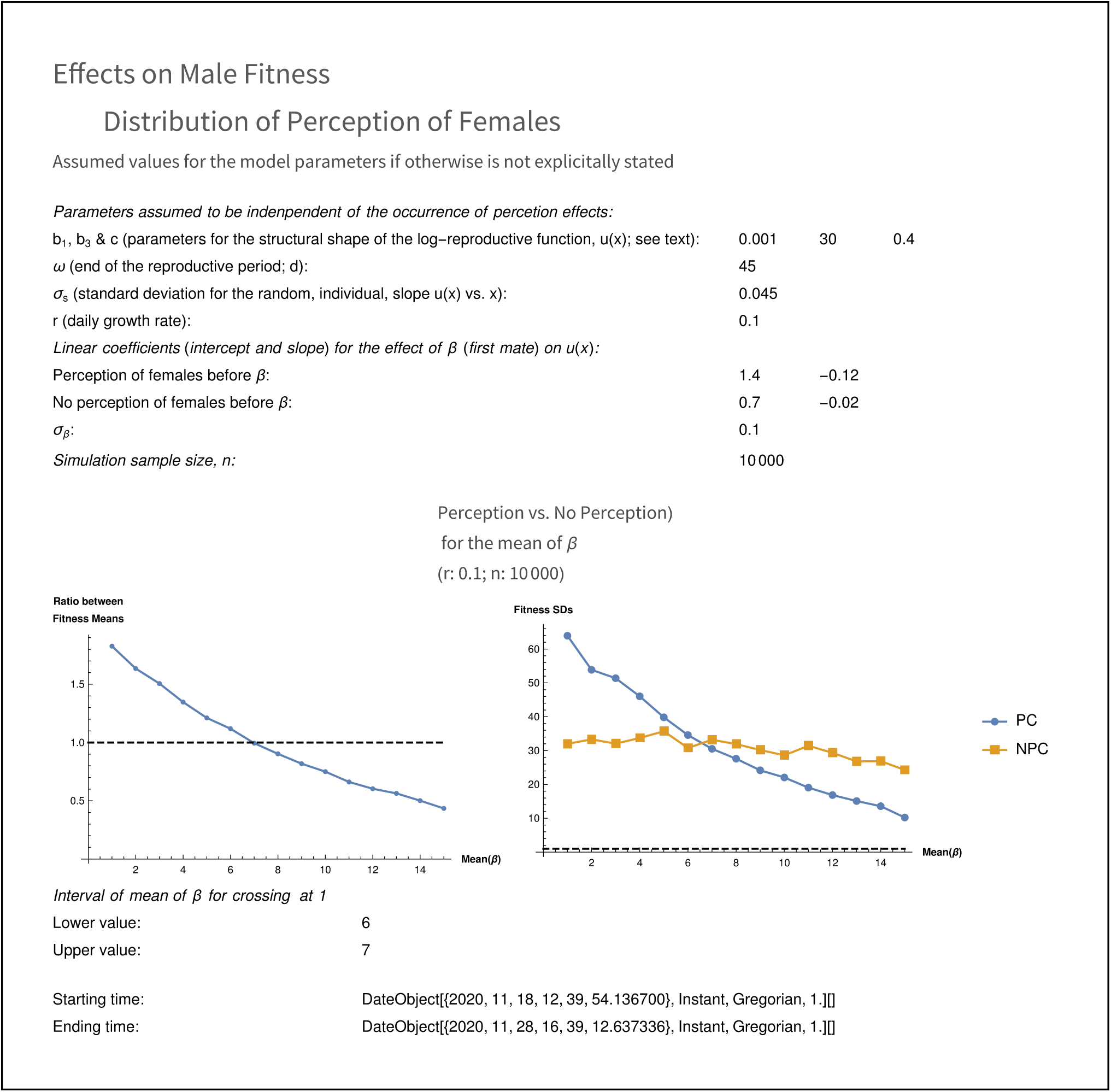
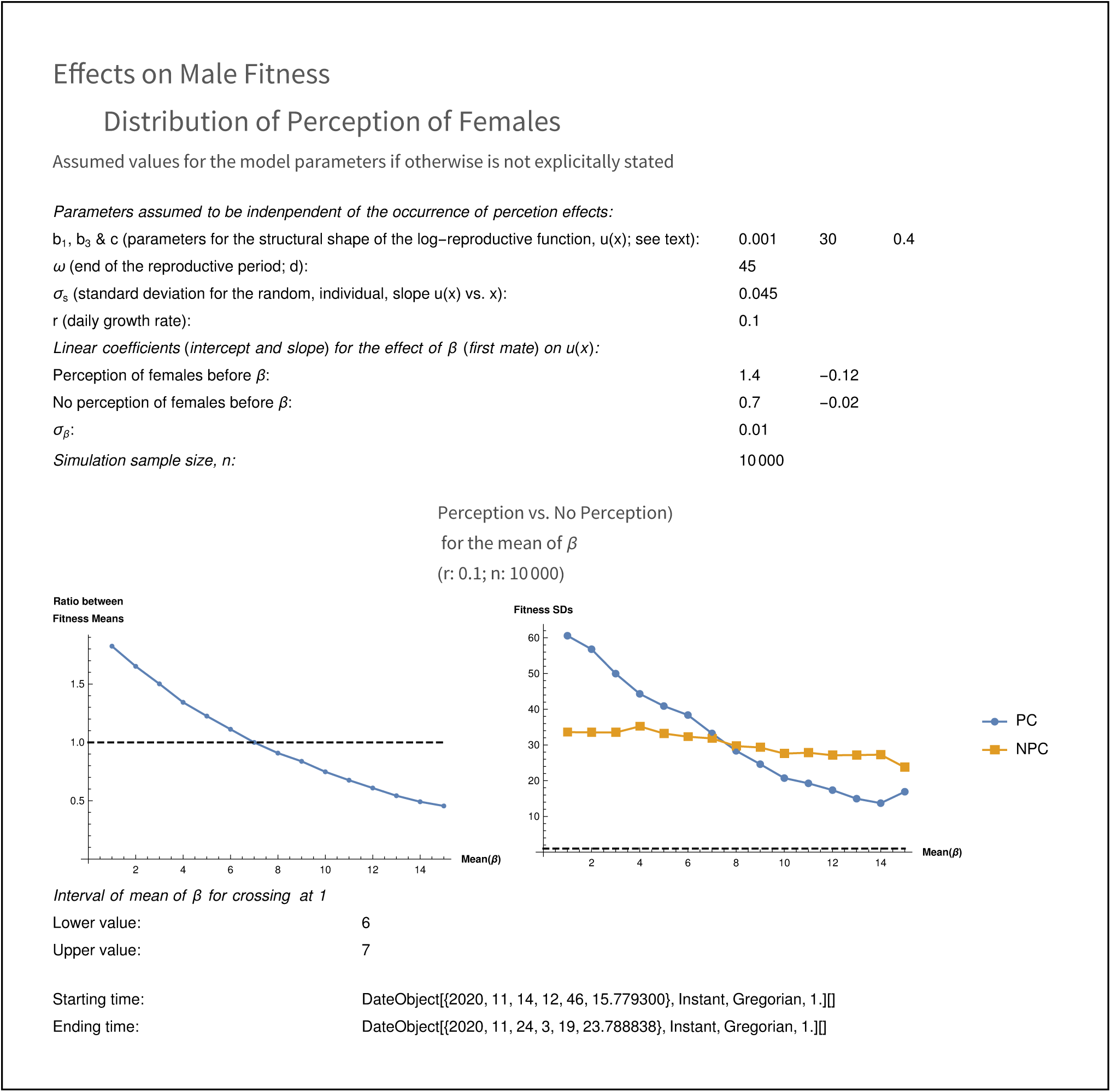
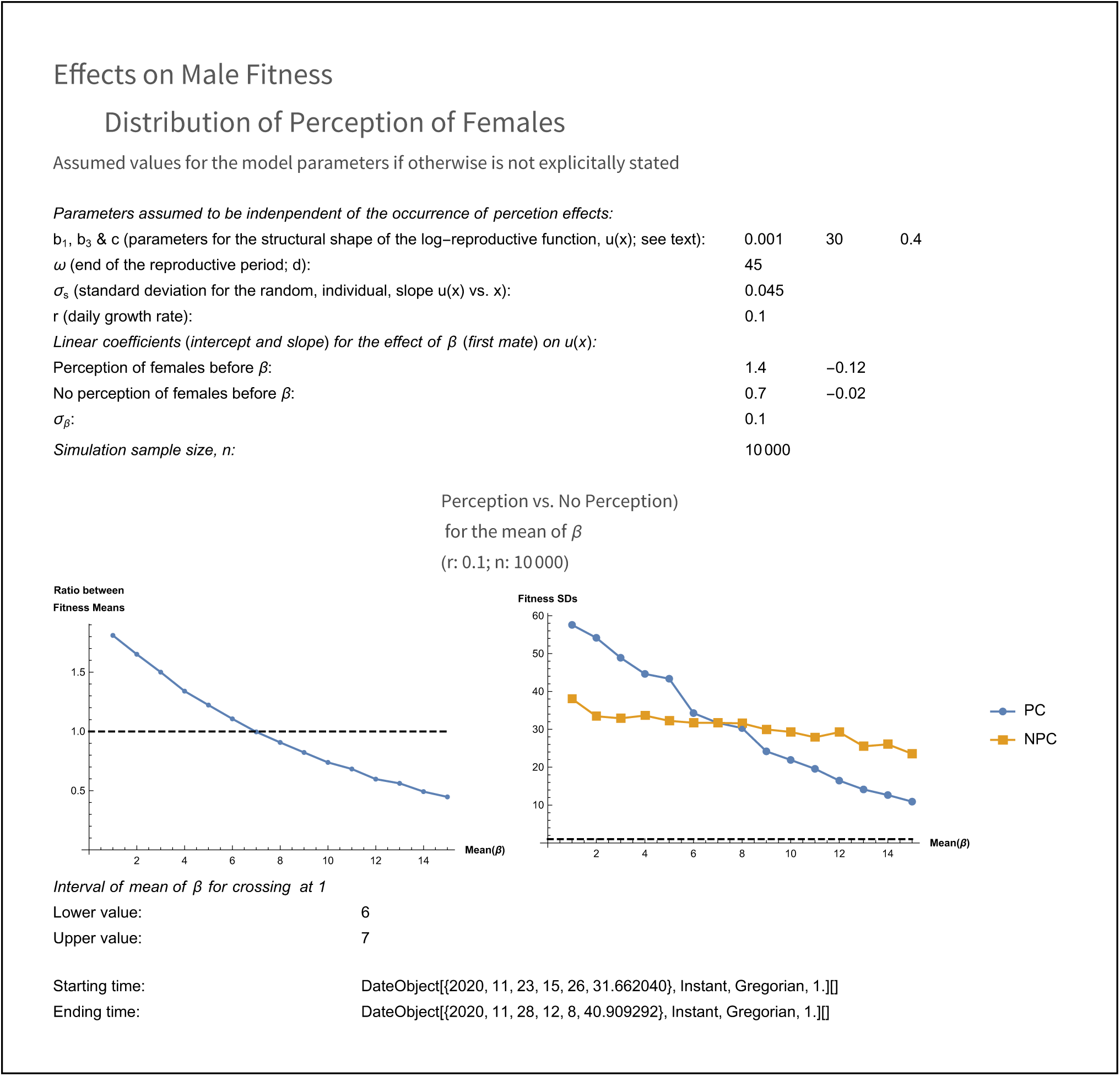
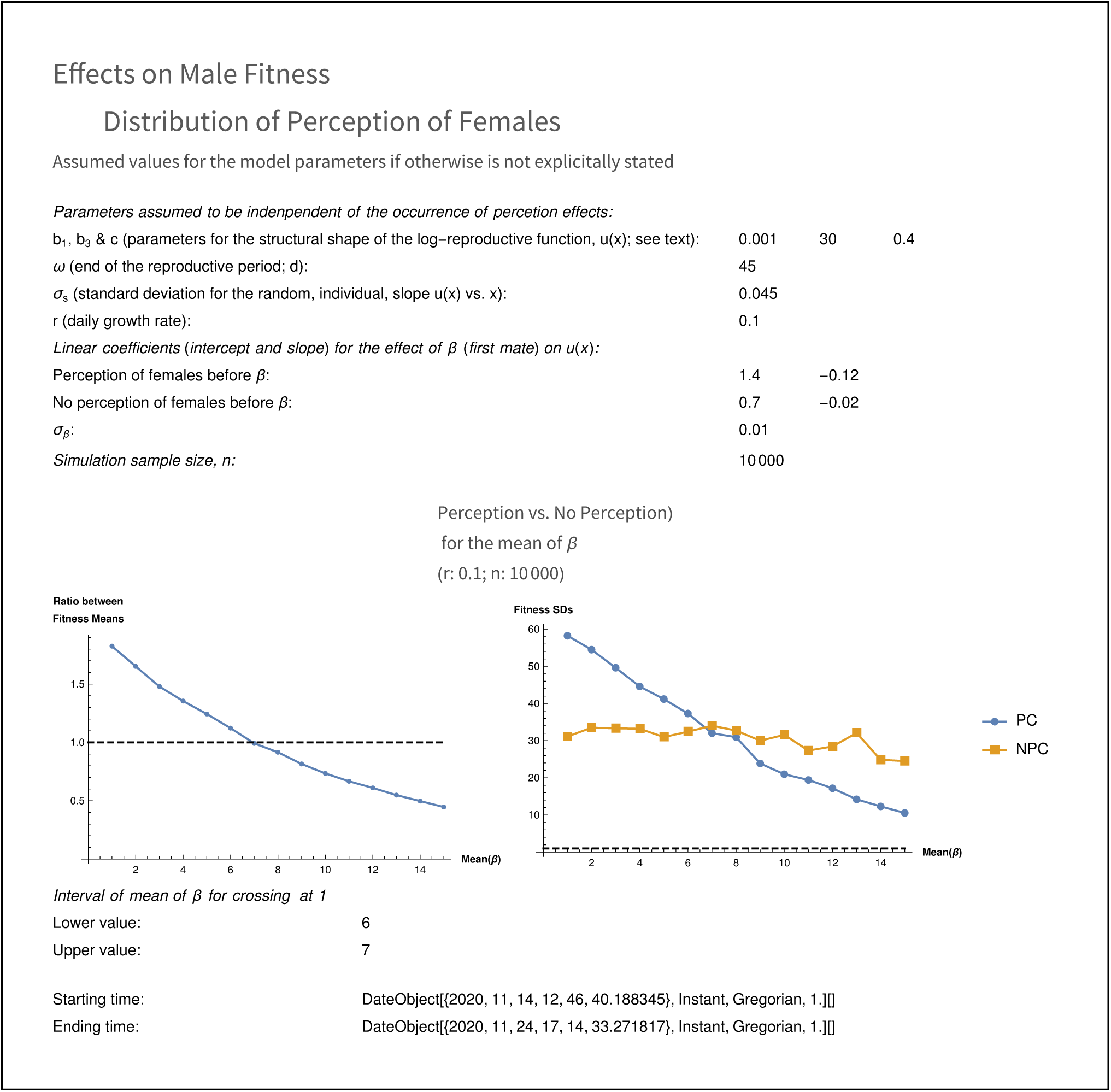
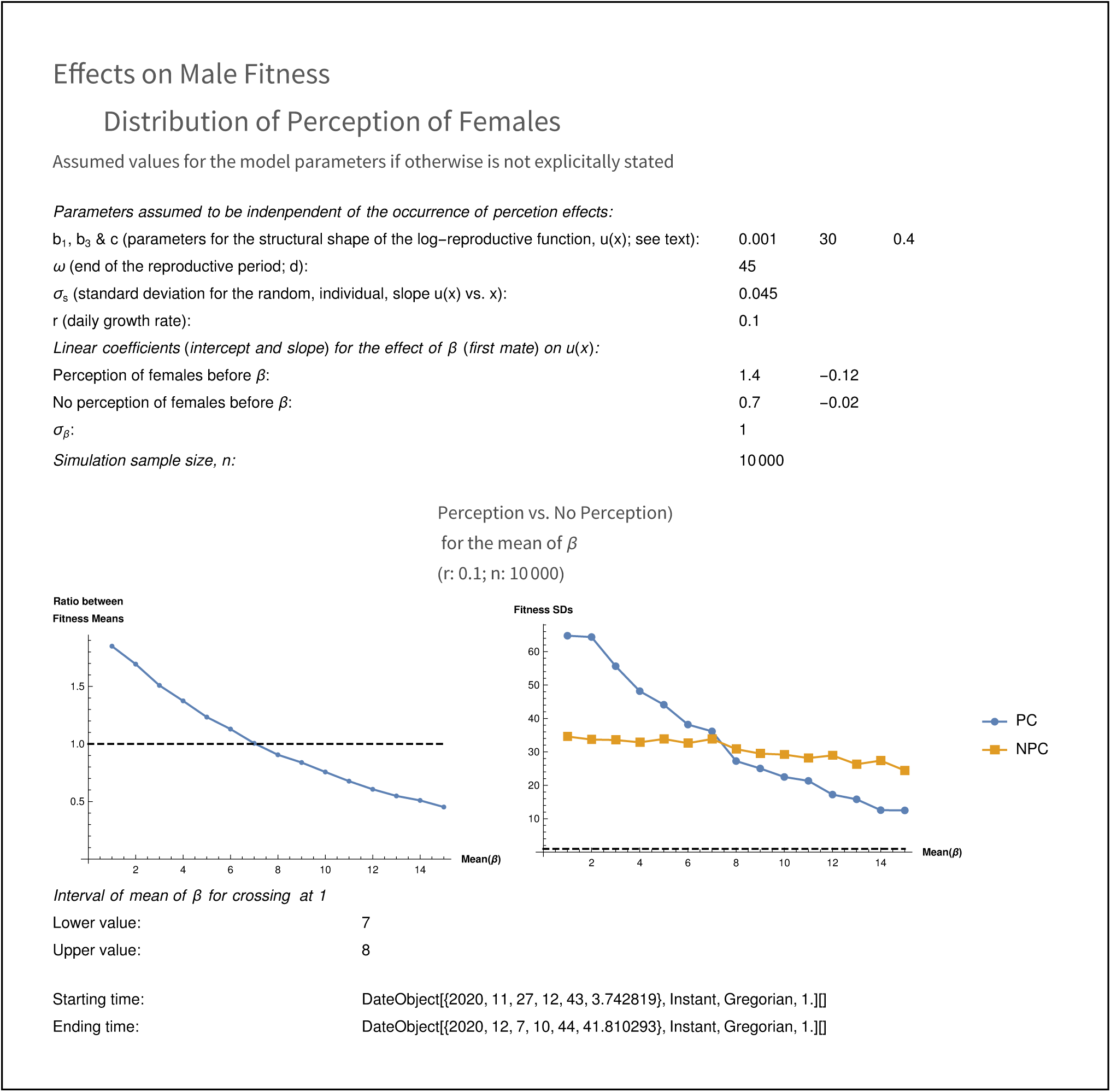
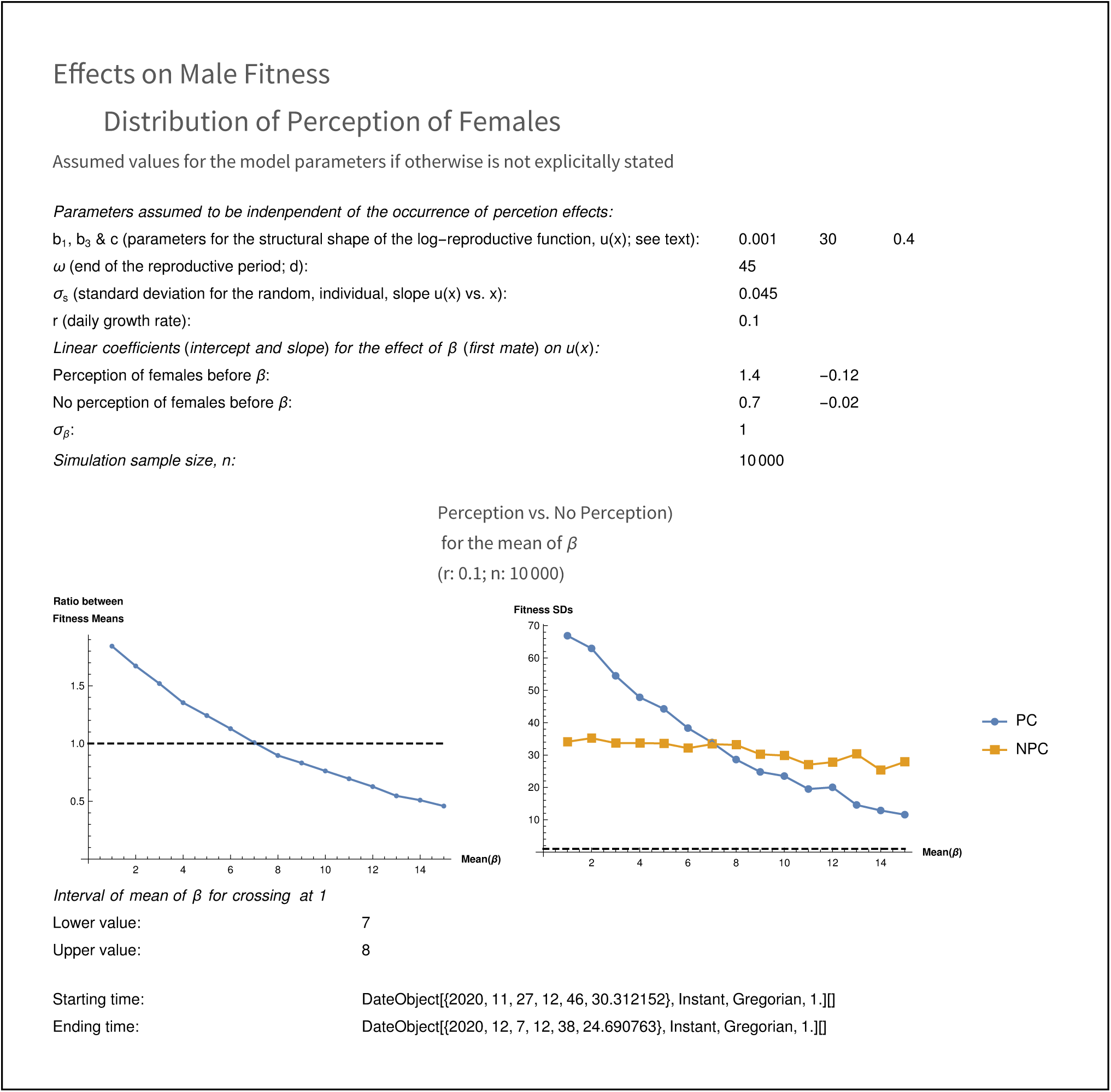
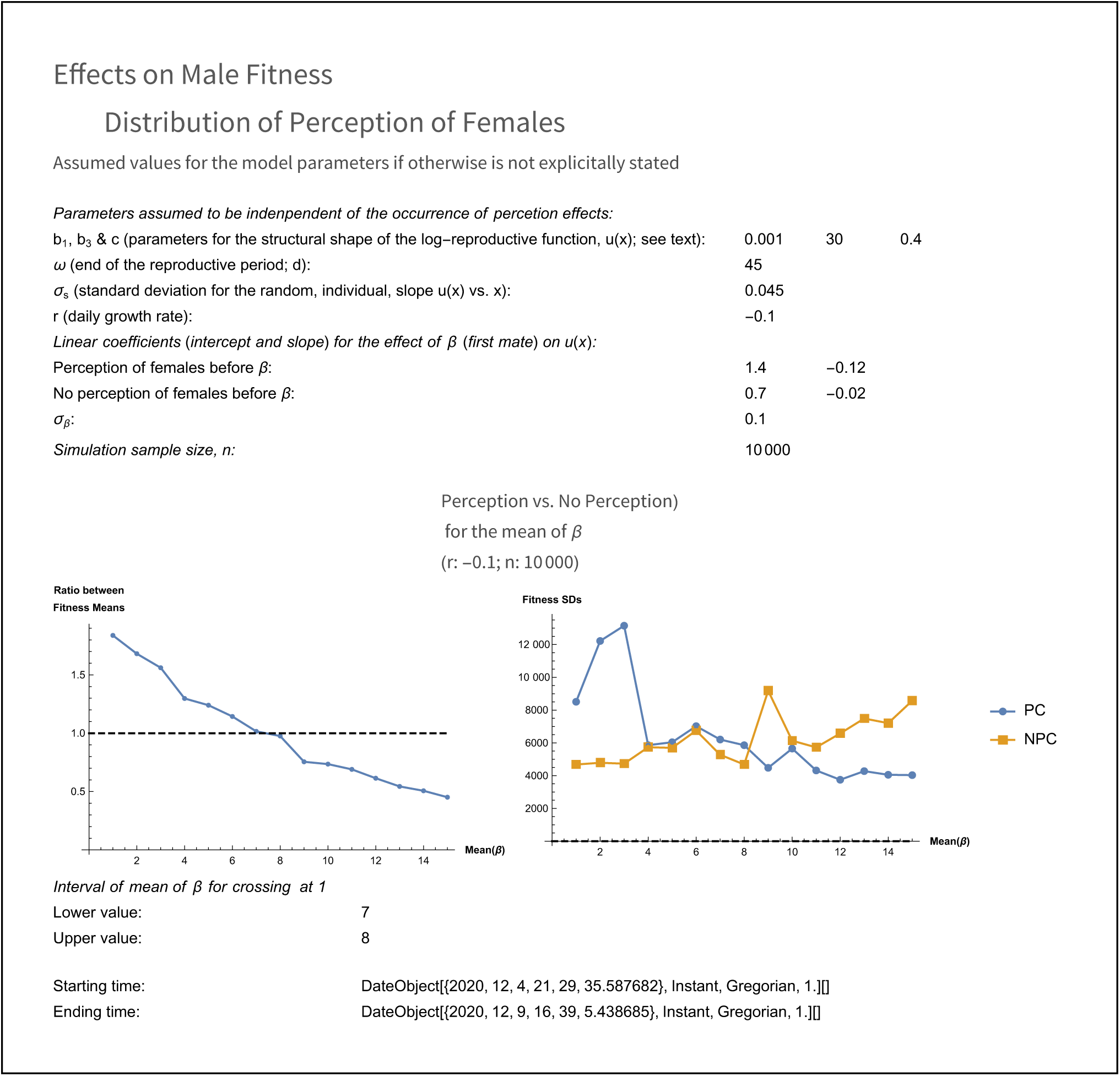
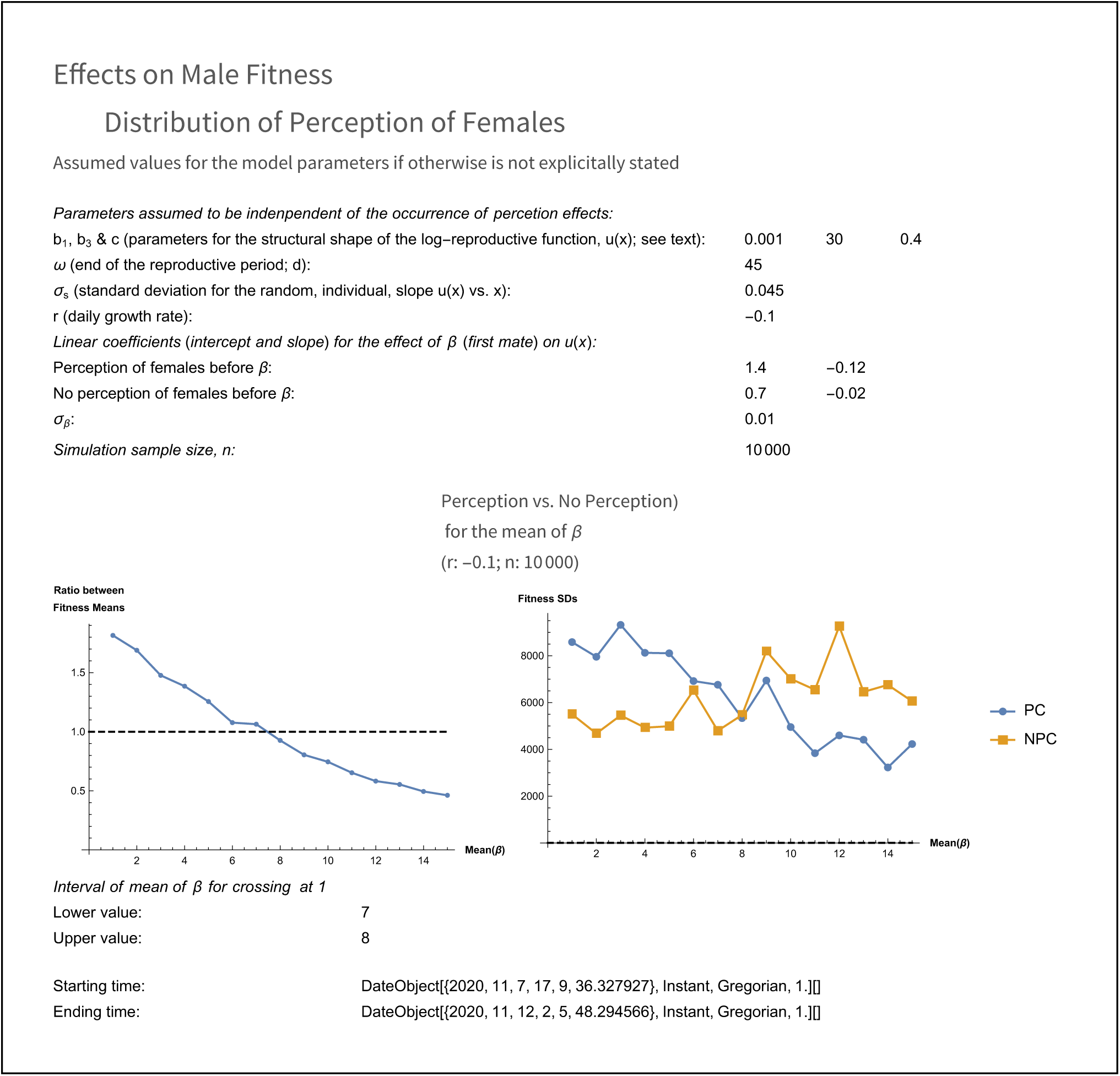
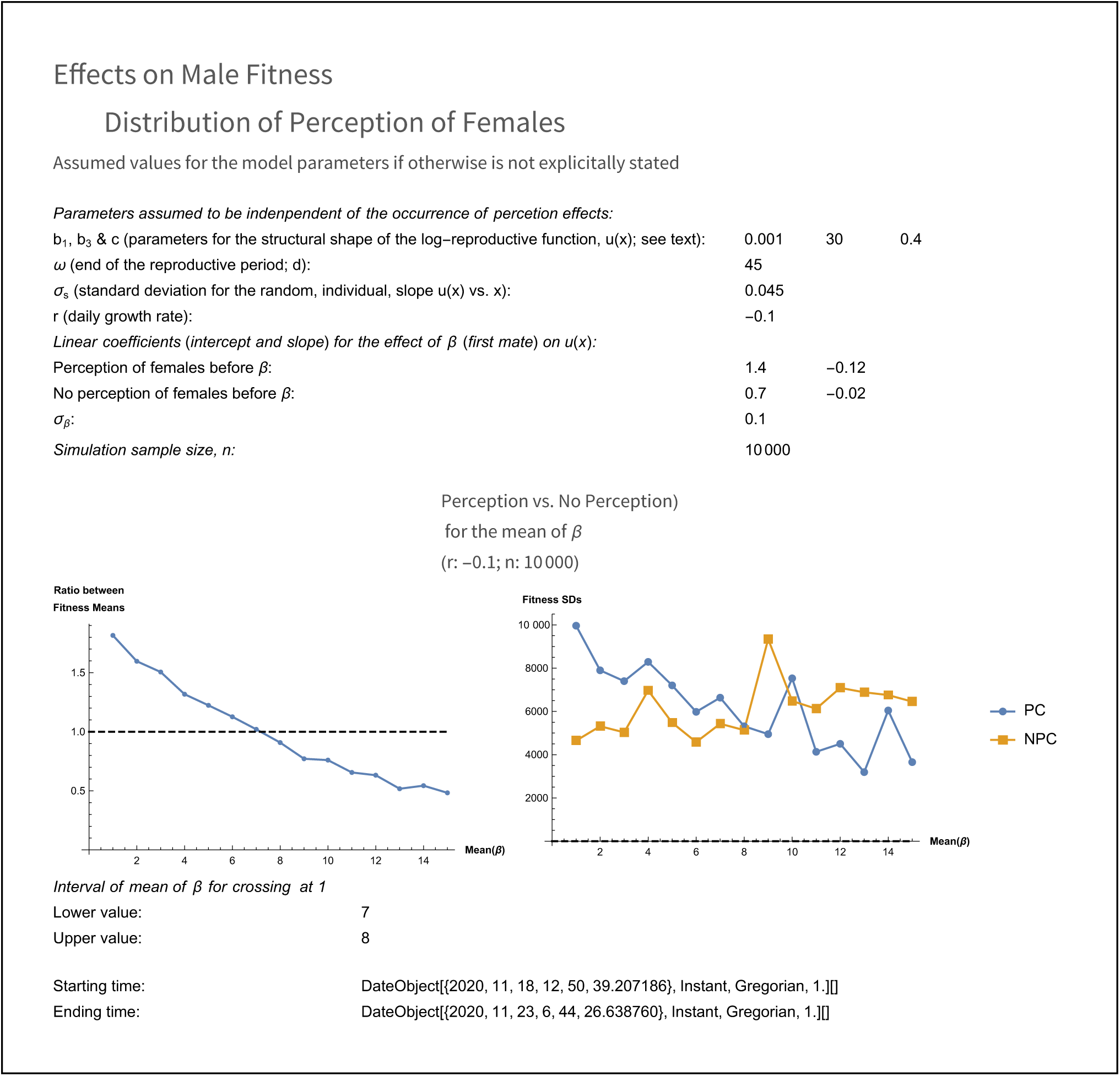
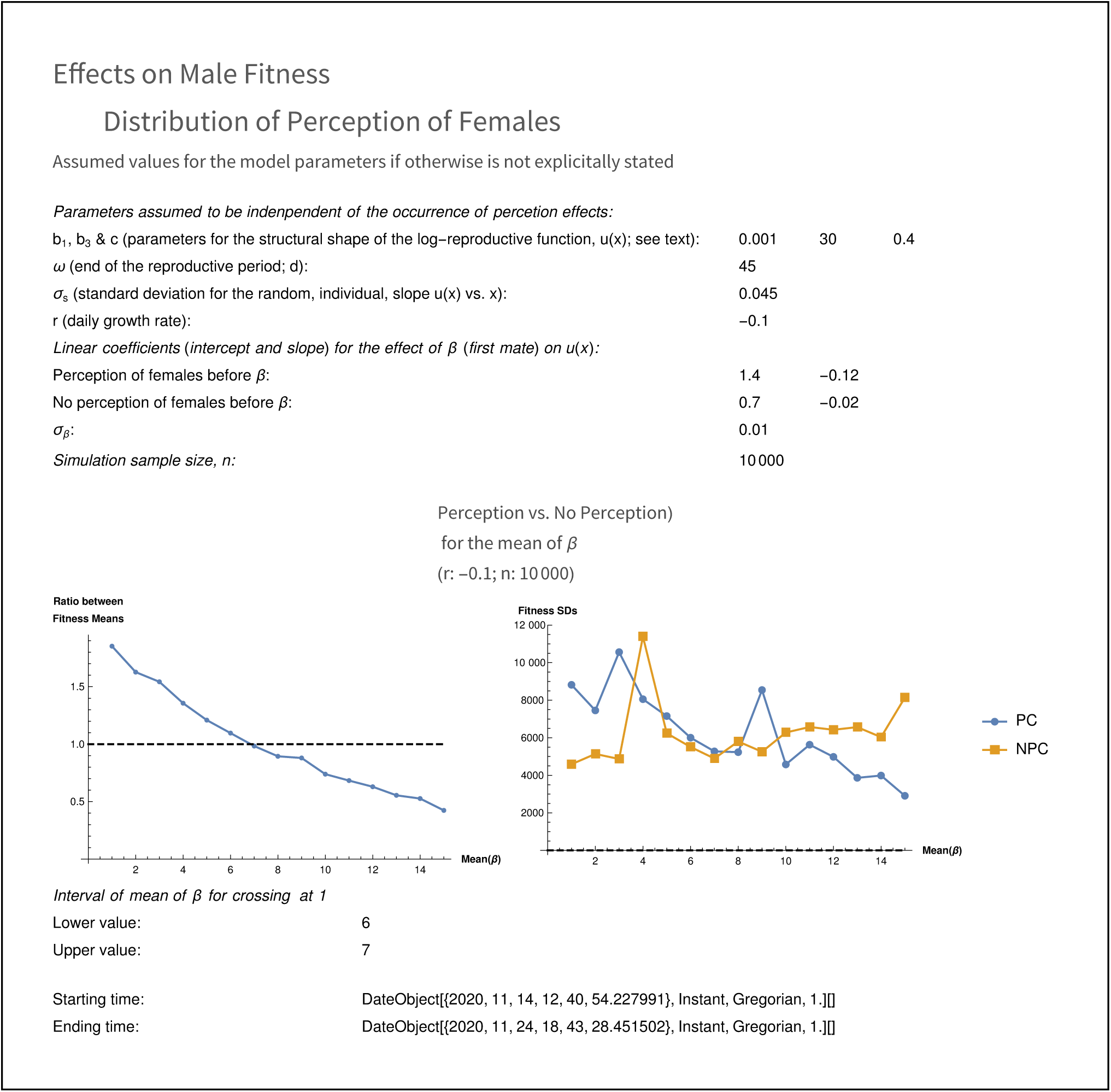
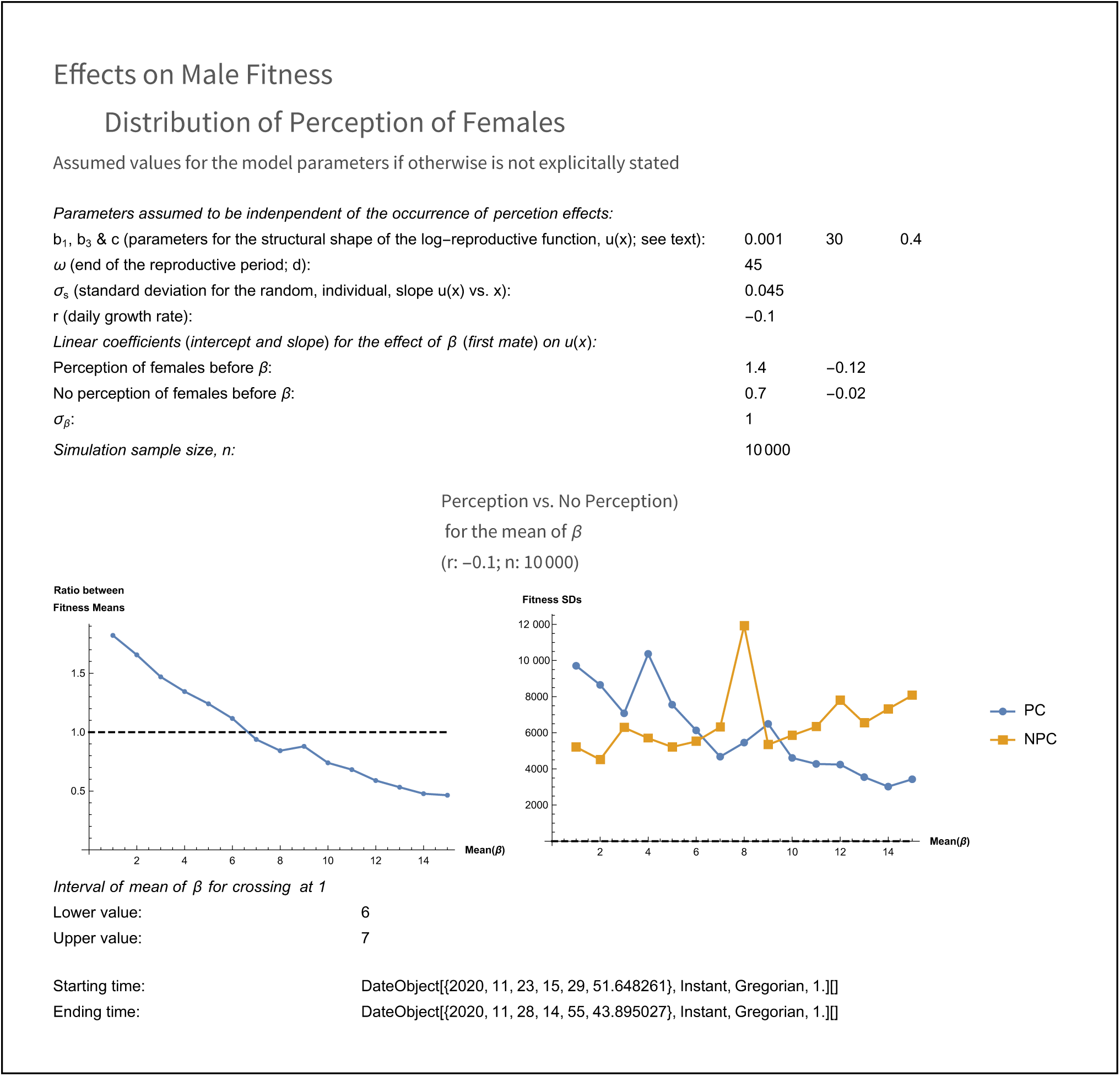
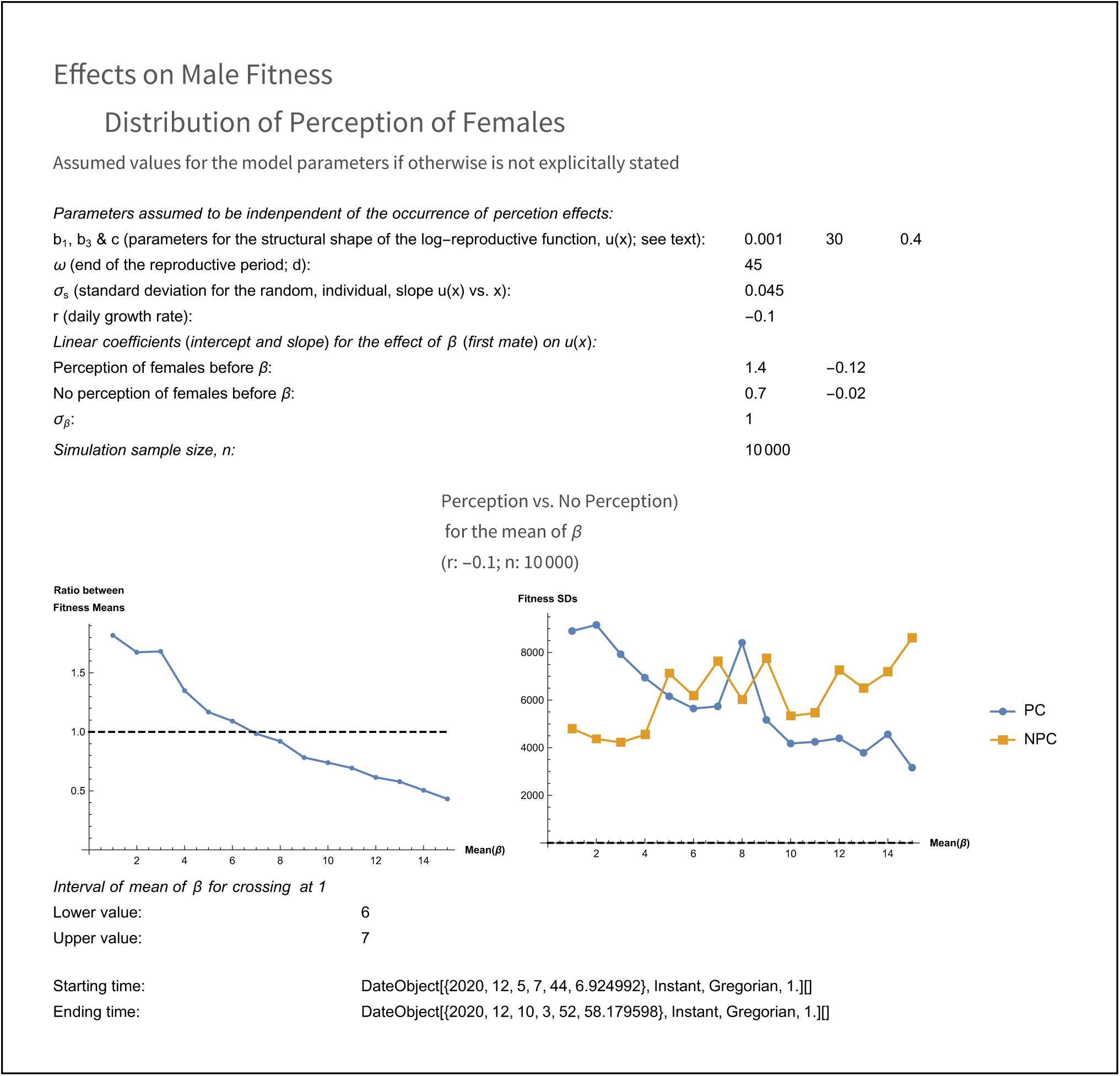
Simulations output across daily population growth rate (r: −0.1, 0, 0.1), mean value of β (from 1 to 15 d), and standard deviation of β (σ_β_: 0.01, 0.1, 1d).

